# Multivoxel Pattern of Blood Oxygen Level Dependent Activity can be sensitive to stimulus specific fine scale responses

**DOI:** 10.1101/798306

**Authors:** Luca Vizioli, Federico De Martino, Lucy S Petro, Daniel Kersten, Kamil Ugurbil, Essa Yacoub, Lars Muckli

## Abstract

At ultra-high field, fMRI voxels can span the sub-millimeter range, allowing the recording of blood oxygenation level dependent (BOLD) responses at the level of fundamental units of neural computation, such as cortical columns and layers. This sub-millimeter resolution, however, is only nominal in nature as a number of factors limit the spatial acuity of functional voxels. Multivoxel Pattern Analysis (MVPA) may provide a means to detect information at finer spatial scales that may otherwise not be visible at the single voxel level due to limitations in sensitivity and specificity. Here, we evaluate the spatial scale of stimuli specific BOLD responses in multivoxel patterns exploited by linear Support Vector Machine, Linear Discriminant Analysis and Naïve Bayesian classifiers across cortical depths in V1. To this end, we artificially misaligned the testing relative to the training portion of the data in increasing spatial steps, then investigated the breakdown of the classifiers’ performances. A one voxel shift led to a significant decrease in decoding accuracy (p<.05) across all cortical depths, indicating that stimulus specific responses in a multivoxel pattern of BOLD activity exploited by multivariate decoders can be as precise as the nominal resolution of single voxels (here .8 mm isotropic). Our results further indicate that large draining vessels, prominently residing in proximity of the pial surface, do not, in this case, hinder the ability of MVPA to exploit fine scale patterns of BOLD signals. We argue that tailored analytical approaches can help overcoming limitations in high-resolution fMRI and permit studying the mesoscale organization of the human brain with higher sensitivities.

## 2. Introduction

Largely due to the ability to achieve relatively high spatial and temporal resolution functional images simultaneously across the whole brain, functional magnetic resonance imaging (fMRI) has become one of the most powerful tools to study the human brain non-invasively over the last 25 years. At ultra-high field (UF, 7 Tesla and above), functional voxels span the sub-millimeter range, measuring 0.8 mm isotropic (e.g. Muckli et al., 2015; for review see Lawrence et al., 2017), 0.65 mm isotropic over small regions (Heidemann et al., 2012); or even 0.45 mm using super resolution techniques (e.g. Vu et al., 2018). These high-resolution images allow the recording of blood oxygenation level dependent (BOLD, Ogawa et al., 1992) responses at the level of cortical layers and columns. UF fMRI therefore provides the unique opportunity to investigate the organizing principles of the human cortex at the mesoscale level, narrowing the gap between invasive animal electrophysiology and human neuroimaging (De Martino et al., 2018).

However, this sub-millimeter resolution is only nominal in nature, because a number of factors limit the point spread function of gradient echo (GE) BOLD responses and, ultimately, the sensitivity to fine-grained functional structures. These factors include voxel blurring along the phase encoding direction and proximity to large draining blood vessels. Studies investigating the point spread function of GE BOLD responses at 7 T have shown that it spreads beyond the millimeter range, with an upper limit of approximately 2 mm (Shmuel et al., 2007; Uludağ and Blinder, 2017). More recently though, it has been argued that these estimates fail to account for the spread of the neuronal response as it relates to the size of receptive fields and their scatteredness in V1 (Chaimow et al., 2018). Chaimow et al. (2018) suggest that when minimizing the contribution of macroscopic vessels, the point spread at 7 T for GE BOLD acquisitions is closer to 1mm than 2mm, approaching the nominal resolution of single voxels (in this case 0.8 mm isotropic).

Moreover, large veins also significantly modulate BOLD amplitude, leading to an increase in signal towards the pial surface, especially for GE recordings (e.g. Goense and Logothetis, 2006; Goense et al., 2007; Ress et al., 2007; Polimeni et al., 2010; Koopmans et al., 2010; Koopmans et al., 2011).

The implementation of appropriate analytical strategies may help to circumvent the impact of large draining vessels on biasing BOLD signal responses to outer cortical layers. For example, differential mapping techniques, along with the presence of pseudo-periodic functional structures (Yacoub et al. 2007), can permit the mapping of orientation preference columns with high field spin echo (SE) fMRI, despite the limited spatial resolution and/or functional precision.

For the highly desirable GE BOLD signal, however, it remains to be determined whether and how neuroscientists can fully exploit the high spatial resolution data achievable at UF, in order to investigate functional profiles of human cortical columns and layers. To this end, analyzing the information contained in voxel populations using multivoxel pattern analysis (MVPA) as opposed to average response amplitudes may represent an appealing analytical strategy to maximize fMRI sensitivity to fine-grained cortical features (Kriegeskorte et al., 2007). Kamitani and Tong (2005) employed MVPA to successfully decode orientation tuning in human V1 at 3 Tesla, with a voxel resolution that spanned well beyond the millimeter range (see also Haynes and Rees, 2005). With the promise of retrieving information that would otherwise remain inaccessible, MVPA, such as linear support vector machines (SVM), have become widely used.

This account has been challenged recently, and the nature and spatial scale of the information exploited by the MVPA called into question. A number of studies have argued that orientation decoding may rely on coarser global maps that co-vary with micro-scale features (e.g. Sasaki et al., 2006; Mannion et al., 2010; Freeman et al., 2011; Op de Beeck, 2010). An example of such a coarse-scale organization that could account for orientation decoding in V1 is radial-preference retinotopic maps (Sasaki et al., 2006). The somewhat unresolved debate sparked by these opposing views has motivated several neuroimaging studies to assess whether MVPA is sensitive to fine or coarse spatial patterns of multivoxel BOLD activity (e.g. Mannion et al., 2010; Seymour et al., 2010; Alink et al., 2013; Op de Beeck, 2010; Freeman et al., 2011; Chaimow et al., 2011), with the possibility that orientation decoding could be underpinned by both coarse as well as fine scale structures.

With the growing availability of UF scanners, the question of whether MVPA decoding relies on fine-grained spatial information becomes topical for the neuroimaging community. As mentioned above, GE BOLD is limited in spatial specificity, casting doubts on the spatial integrity of sub-millimeter fMRI maps. However, a demonstration that, unlike univariate amplitudes, MVPA effectively exploits finer grained information from GE-BOLD data, taking full advantage of the sub-millimeter resolutions, stands to increase the utility of high field high resolution GE BOLD. Within the context of this paper, we define univariate BOLD as the average BOLD response (or the contrast of the average BOLD responses elicited by 2 conditions) across all voxels within a given region of interest (ROI).

In this work, we re-analyzed feedforward and feedback cortical depth dependent data previously acquired 7 T GE EPI with 0.8 mm isotropic functional voxels (Muckli et al., 2015, Figure 1) to determine the spatial scale of stimulus specific responses to which MVPA is sensitive.

**Figure 1.**
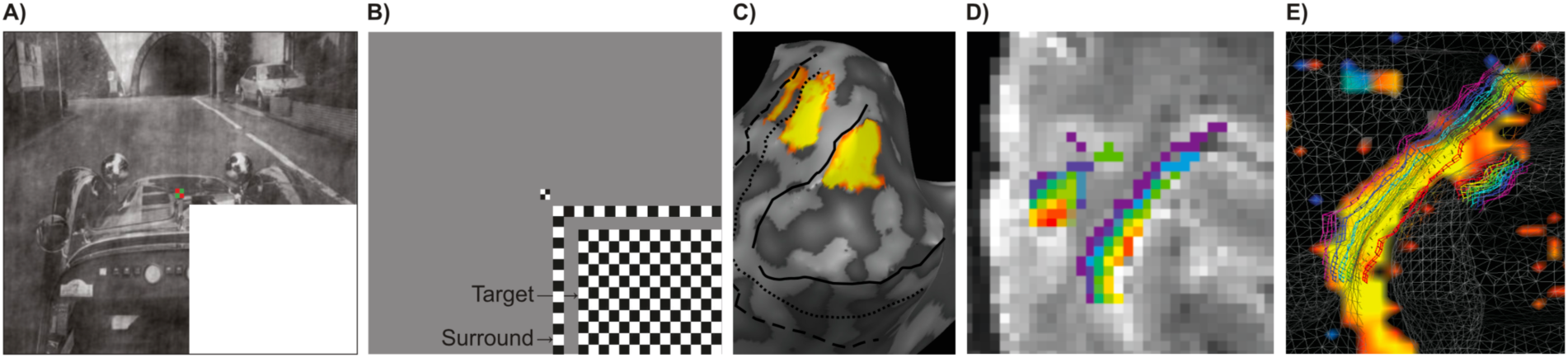
(A) Example stimulus for the ‘‘feedback’’ condition, in which the lower right quadrant was occluded by a white mask. The ‘‘feedforward’’ condition comprised the full image (not shown). (B) ‘‘Target’’ and ‘‘surround’’ checkerboards (presented individually during scanning) to locate voxels responding to the lower right visual field. (C) Left hemisphere inflated cortical reconstruction of a subject, overlaid with a contrast of target response greater than surround response in V1, V2 and V3. (D) Example of cortical depths (purple, close to white matter and red, close to the pial surface) overlaid onto GE-EPI images. (E) White transparent wireframe shows cortical surface reconstruction along the grey-white matter boundary with the superimposed iso-surface cortical depth grids (inner in purple, to outer in red) of a representative subject’s left occipital cortex (sagittal view). The visual activation map results from the contrast t_target_ > t_surround_ ∩ t_target_ > 0 ∩ t_full-stimulus_ > t_occluded-stimulus_. Figure reproduced and adjusted with permission from (Muckli et al. 2015).

To this end, we artificially misaligned the pattern structure in increasing spatial steps and investigated the breakdown of the classifier performance. We reasoned that if the spatial scale of stimulus specific responses to which decoding is sensitive is that of a single voxel, even a 1 voxel misalignment should lead to a significant decrease in decoding accuracy. To test this hypothesis, we first ran a simulation on synthetic data generated with realistic signal-to-noise (SNR) properties.

We then tested the performance of 3 classifiers: linear SVM; Linear Discriminant Analysis (LDA); and Naïve Bayes Classifier (NBC) on real data. Moreover, to gain insights into whether venous-related amplitude increases also affect the magnitude of univariate differences and decoding accuracy, we directly assessed the relationship of these measures and univariate BOLD across cortical depths by means of correlational analyses.

## 3. Methods

### 3.1 Subjects

We reused data acquired from 4 subjects at the Center for Magnetic Resonance Research (CMRR, Minneapolis, MN, USA) and described in Muckli et al. (2015). All subjects were healthy volunteers with normal or corrected visual acuity. Subjects gave written informed consent and received financial compensation for their participation. The institutional review board for human subject research at the University of Minnesota approved the study.

### 3.2 Stimuli

Stimuli are described in the original study (see Muckli et al., 2015). In summary, subjects viewed three visual scenes (‘car on street’, ‘people at market’, ‘ship in harbor’, Figure 1). We controlled the images for global luminance, contrast and energy, using matlab shine-toolbox (Willenbockel et al., 2010). Scenes were presented in full (‘feedforward’ condition) or masked with an occluder over the lower right visual field (‘feedback’ condition). We presented a set of contrast-reversing checkerboard mapping stimuli for ‘target’ and ‘surround’ regions in each run, and in a separate localizer run. The surround checkerboards mapped the outer 2 degrees of the white occluder and the target mapped the remaining inner section of the occluder (Figure 1). The design of the experiment was comparable to our previous study (Smith and Muckli, 2010), but with the visual stimuli reduced in size by 20% to fit the smaller MRI bore due to the use of a head gradient insert (see below). We kept the width of the ‘surround’ mapping stimulus at 1 degree of visual angle and added an additional 1-degree border between the ‘surround’ stimulus and the edge of the ‘target’ stimulus region. We conducted a separate phase-encoded retinotopic mapping experiment (Petro et al., 2013; Sereno et al., 1995; Muckli and Petro, 2013). Stimuli consisted of a wedge-shaped (22.5 degrees) checkerboard rotating slowly (64s for full 360-degree rotation) around the fixation point in the middle of the screen. A white ‘spider web’ configuration was presented in the background to stabilize fixation together with a center fixation color change task (Schira et al., 2009).

### 3.4 Paradigm

As described in Muckli et al. (2015), the experiment comprised four functional runs of 350 volumes each. An experimental condition was presented for 6 volumes (12s), and each of 6 experimental conditions was presented in a randomized order within a block followed by 12 volumes (24s) of baseline (6×12s+24=96s per block). Mapping blocks consisting of 2 conditions (‘target’, ‘surround’) were presented for 6 volumes (12s) interleaved with 6 volumes (12s) of baseline in between conditions and 12 volumes (24s) at the end of the block (2×12s+12s+24s=60s). A functional run consisted of 6 experimental blocks and 2 mapping blocks and an additional baseline of 2 volumes (4s) at the start of the run (6×96s + 2×48s + 4s = 700s = 350 volumes). Therefore, each experimental condition was presented 24 times across four runs.

The retinotopic mapping run comprised of 12 repetitions of a full rotation lasting 32 volumes (64s), with an extended baseline of 10 volumes (20s) at the beginning and 12 volumes (24s) at the end of the run (resulting in 406 volumes: 12×64s+20s+24s=812s). An additional localizer run comprised 12 repetitions of ‘target’ and 12 repetitions of ‘surround’ mapping, with 25 baseline periods in between, all of which lasted for 6 volumes (12s), resulting in 294 volumes ((12+12+25)x12s=588s).

Subjects viewed the visual stimuli on a projection screen mounted to the rear end of the head coil using a head-coil mounted mirror. A video projector combined with a mirror projected the stimuli onto the screen. Stimuli were presented using Presentation software (Neurobehavioral Systems, CA, USA) for the experiments, and for retinotopic mapping with StimulGL (custom-built stimulation software, Maastricht University, Maastricht, NL). We instructed the subjects to keep fixation to the center of the screen and to perform a color-change detection task at the center of the screen, during both the experimental runs and retinotopic mapping.

### 3.5 MRI Acquisition

MRI data for the first experiment was conducted on an ultra-high magnetic field (7 Tesla, 90cm bore, Magnex Scientific, Abingdon, UK) at the CMRR in Minneapolis (MN, USA). The scanner was driven with a Siemens console (Erlangen, Germany) and used a head gradient insert with a 6-channel receive (1 Tx) array RF coil that covered only the visual areas.

Functional scans were recorded using GE-EPI at high resolution (nominal resolution, isotropic 0.8 mm^3^, TE = 17ms, maximum flip angle (determined by a flip angle map) = 85°, slices = 38, TR = 2000ms, FOV = 128 x 128 mm^2^, matrix: 160 x 160, IPAT = 2, partial Fourier = 6/8, pixel bandwidth = 1375 Hz/pixel). Anatomical scans were acquired at 1mm^3^ using a MPRAGE sequence optimized for T1-weighted (3D MPRAGE) and proton density (PD)-weighted contrast (176 slices, FOV = 136×256 mm2, matrix = 1366×56, voxel size = 1×1×1 mm3).

### 3.6 Anatomical data analysis - cortical depth sampling

All data were analyzed with BrainVoyager QX 2.8. Proton density scans with identical slice positioning were used to remove spatial intensity inhomogeneities from T1 scans by dividing the T1 by the PD images (Van de Moortele et al., 2009). We manually adjusted inner and outer grey matter boundaries along the local intensity values to eliminate pial blood vessels and to correct for GE-EPI distortions. We used relative cortical depth values to create Laplace-based equipotential grid-lines (i.e. solving the Laplace equation to obtain a smooth vector field and then create smooth meshes directly within the grey matter boundaries – e.g. Kemper et al., 2018) at six depths (from inner to outer 90%, 74%, 58%, 42%, 26%, and 10% depths, Figure 1). The gridlines were calculated smoothly at a highly up-sampled spatial coordinate system (De Martino et al., 2014). In a subsequent step, we used smooth gridlines to assign voxels to a respective cortical depth. Individual voxels were allowed to belong to adjacent depths. The depth gridlines covered the cortical representation of the occluded image section in the lower right visual field quadrant of retinotopic area V1d (Figure 1). We saved the layered regions of interest as BrainVoyager QX VOI files (volume of interest).

### 3.7 Functional data analysis

We pre-processed the fMRI data using slice scan timing corrections (sinc interpolation), 3D rigid body motion correction (sinc interpolation), intra-session alignment to the functional data of the last run, and temporal high pass filtering of 4 cycles. We aligned functional data to anatomical data with manual adjustments and iterative optimizations. We used activation maps of retinotopic mapping to optimize segmentation and alignment. Specifically, after T1 based segmentation, the grey matter boarders were projected into EPI space and locally optimized according to the mean EPI image, which provides additional information regarding the spatial location of grey and white matter boundaries and outer pial surface. To further asses the quality of alignment and segmentation, we projected the activation maps from the retinotopic mapping runs to the segmented cortical ribbon. We visually inspected the quality of alignment and segmentation and optimized either or both accordingly. We implemented this procedure on the assumption that activity originates in the grey matter.

Analysis of functional data included general linear model (GLM) estimation of averaged conditions and single trials. We generated design matrices by the convolution of a double gamma function with a “boxcar” function (representing onset and offset of the image stimuli).

Independently per voxel and functional run, we implemented a classic general linear model (GLM) analysis (least squared minimization stress) to estimate the activation triggered by each single image block (i.e. single trial estimation modeling). We computed design matrices by convolving a double gamma function with a “boxcar” function (representing onset and offset of the image stimuli). Each design matrix thus consisted of 350 rows, representing the runs’ temporal dimension (i.e. volumes), and 41 columns, one per each visual stimulation trial plus the intercept term. Of 40 visual trials, 36 represented our experimental conditions: 6 trials x 2 experimental conditions (i.e. full and occluded images) x 3 visual scenes; and 4 represented the mapping stimulus: 2 trials x 2 mapping conditions (i.e. ‘target’ and ‘surround’ checkerboard stimuli). Methods up to this point are described in Muckli et al. (2015).

### 3.8 General Decoding procedure

Linear support vector machine (SVM) decoding analysis was performed using SVM algorithms as implemented by the LIBSVM toolbox (Lin, 2011), with default parameters (notably C = 1). Linear discriminant analysis (LDA) was implemented using the “fitcdiscr” and “predict” functions inbuilt in Matlab’s statistic toolbox (Matlab, The Mathworks Inc, 2014). Naïve Bayes classifier was implemented using the function Classify from Matlab’s statistic toolbox with the ‘diaglinear’ option. Note that, before being input to the classifiers, the activity of each voxel was scaled using the same scaling factors for training and testing sets (Lin, 2011). Firstly, the training portion of the data was normalized within a range of −1 to 1. This normalization was achieved by subtracting from the training data set its minimum value (to set the minimum to 0), dividing the resulting data set by its maximum (to scale the data between 0 and 1), multiplying by 2 and subtracting 1 (to scale the data between −1 and 1). The testing portion of the data was then scaled with the same procedure, but using scaling factors obtained from the training portion of the data (Lin, 2011). All decoding analyses were performed only on voxels responding to target more than to surround (defined by the contrast t_target_ > t_surrond_ ∩ t_target_ > 0). We have described the decoding analyses in more detail previously (Smith and Muckli, 2010).

In brief, we trained all classifiers (linear pattern) to map between activation patterns from three scenes (full feedforward images in experiment 1) or between occluded scenes. We tested the trained classifiers on independent data (leave one run out cross validation). We measured the classifier performance of each cortical depth independently and we tested the single trial classification for significance using permutation testing (10000 iterations of randomly assigned labels). To determine the empirical chance level, we implemented the following procedure. Independently per subject cortical depth, signal and misalignment extent, we randomly shuffled the labels of the classifiers’ input prior to the training phase. We then performed training and testing with shuffled labels. We repeated this procedure 10000 times to produce a null distribution of decoding accuracy. We therefore sorted the label shuffled accuracy scores and selected the 95% largest score as the empirical chance level. Statistical significance was inferred when the low confidence interval of the unshuffled decoding accuracy (computed across cross-validated folds) was larger than empirical chance (i.e. p<.05).

### 3.9 Artificial misalignment

To directly measure whether MVPA is capable of relying on stimulus specific fine scale responses, we developed a simple data driven approach that builds upon the impact of misalignment between training and test ROIs on decoding accuracy. Independently per cortical depth, we trained a classifier on the original ROI and tested its accuracy/performance on a number of misaligned sites.

We parametrically *shifted* the test site 0 (i.e. no misalignment) to 5 voxels relative to the training site.

To determine whether the drop in decoding accuracy following misalignment can be directly related to the spatial precision of stimulus specific responses, we generated synthetic sets of data while parametrically varying the precision of 3D patterns of simulated BOLD activity via 3D convolution (Figure 2).

**Figure 2.**
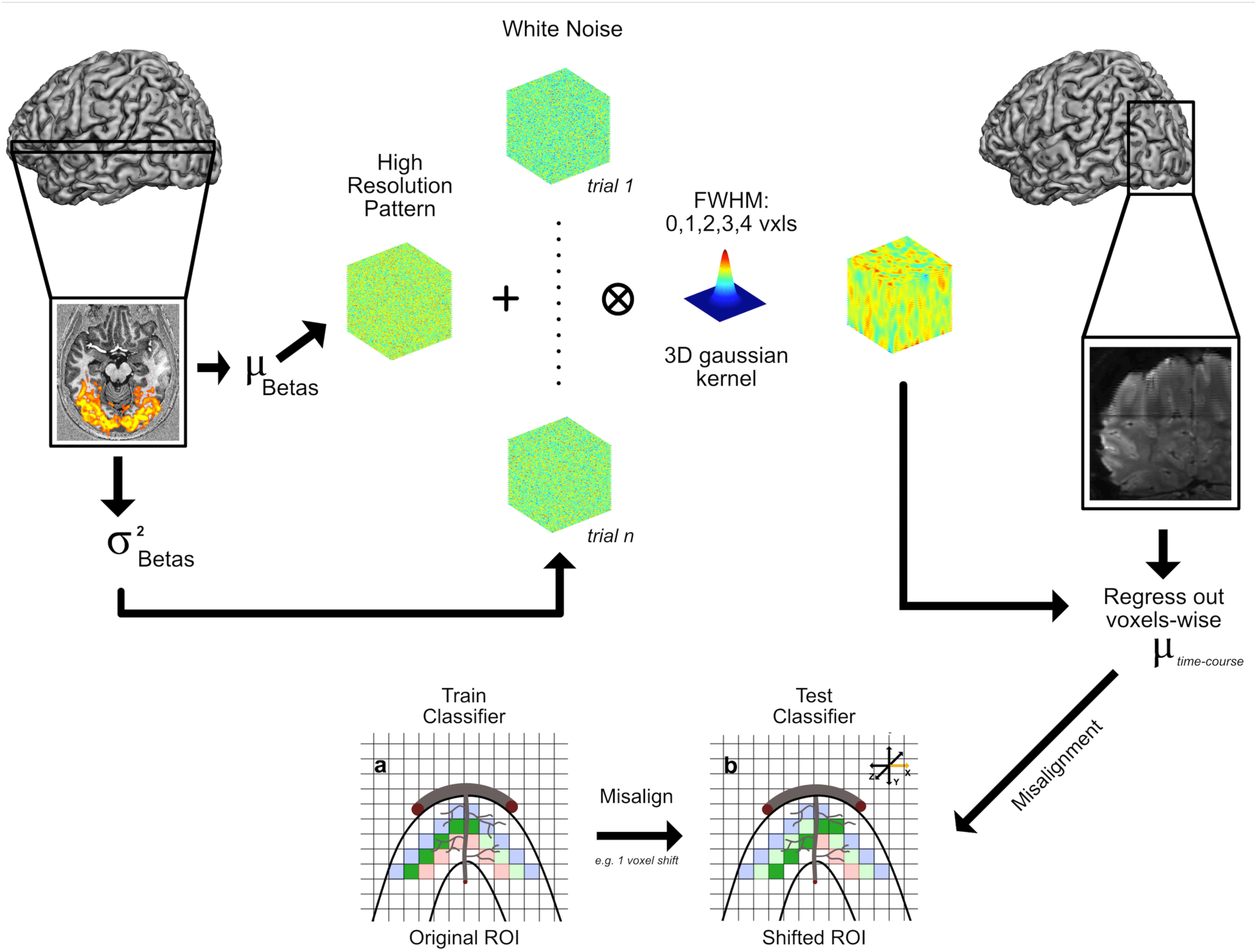
Synthetic data generation procedure.

For each subject and cortical depth, we generated synthetic data mimicking the BOLD activation triggered by 2 visual conditions (see Figure 2). We began by estimating mean betas and variance across runs and trials. We then generated 2 (i.e. one per visual condition) 3D pseudo-random high-resolution patterns of normally distributed white noise with mean of 0 and variance of 1 (using the “randn” function in matlab). These 2 3D textures represent the true (i.e. no noise) multivoxel pattern of our synthetic conditions. We therefore added the previously estimated betas mean to each condition to attain comparable mean activation. Next, we generated patterns of noise. Using a comparable procedure, we proceeded to produce as many normally distributed high-resolution white noise patterns (with mean 0 and variance of 1) as the total number of experimental trials (in this case 48 – i.e. 6 trials x 4 runs x 2 conditions). We therefore used the previously estimated variance to scale the noise patterns so that the variance across the 24 3D noise textures of each image was comparable to that estimated for our real data set. We added each scaled noise texture to the signal patterns, producing 24 trials per condition with the same underlying spatial structure and different amounts of noise. We then smoothed the 3D multivoxel patterns of each simulated trial by convolving it with a 3D Gaussian kernel with a full-width half-max (FWHM) of 0 (i.e. no smoothing), 1, 2, 3 and 4 voxels, to simulate voxel resolutions of .8, 1.6, 2.4, 3.2 and 4 mm isotropic respectively. We finally added the noise and signal texture patterns to our baseline.

This process led to the generation of 2 synthetic multivoxel patterns of betas per simulated voxel resolution with comparable means and distinct spatial patterns of activation, faithfully reproducing the activation profile of our real data. We then carried out our misalignment approach on the synthetic data sets.

We then proceeded to implement this analysis to our data. We used two different, yet complementary, approaches: 1) *volumetric driven* and 2) *surface grid driven* misalignment. While misalignment was performed in volume space in both approaches, unlike the volumetric misalignment, the surface grid driven approach ensured that all spatial offsets of the training ROI were confined within a given cortical depth (Figure 3, see below for more details).

**Figure 3.**
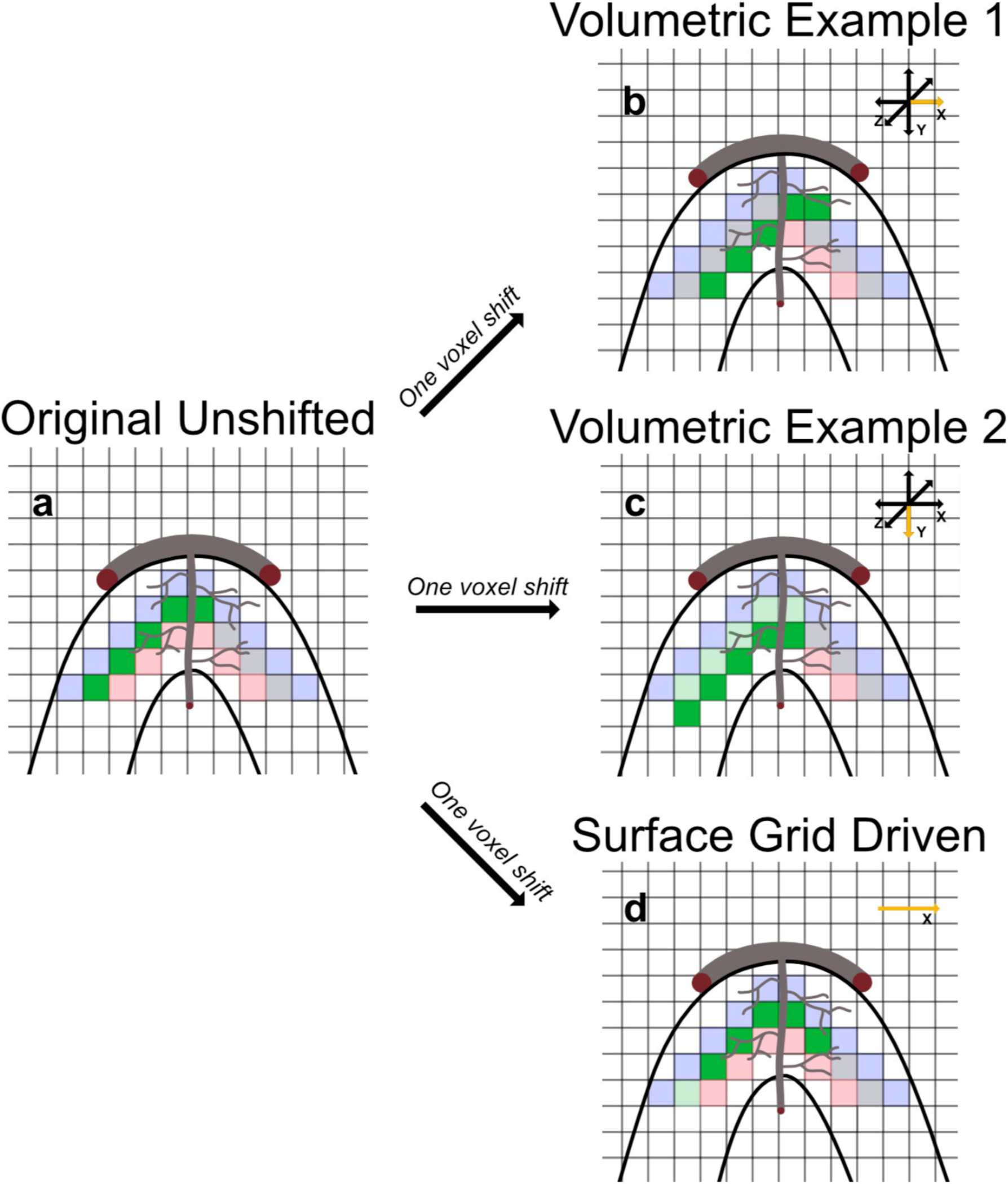
Volumetric and Surface Grid Driven Misalignment. 2-dimensional cartoon representation of the 2 types of artificial misalignments implemented. The curved black lines represent a portion of the cortical sheet. The largest gray curved line, tangential to the cortical sheet, represents a large vessel on the pial surface; while the other curved gray line represents a penetrating draining vessel. The thinner wavy gray lines represent smaller vessels such as capillaries. The pale red, green and blue colored squares symbolize voxels belonging respectively to inner, mid and outer layers, and the solid green squares represent voxels in the middle layers belonging to the retinotopic representation of the occluded bottom right hand quadrant of the visual field (i.e. the ROI that will be misaligned). Note that while we segmented the cortical sheet into 6 layers, this figure, for simplicity, only shows 3 layers. Panel a) depicts the original, unshifted ROI. Panel b) shows the effect of 1 voxel volumetric misalignment. The arrows on the top right corner represent all possible directions along which misalignment occurs. The yellow arrow indicates the dimension along which misalignment has occurred (in this case to the right along the x axis). Panel c) is identical to panel b) apart from the dimension along which the training ROI was misaligned (i.e. downwards along the y axis). Panel d) portrays an example of 1 voxel surface grid driven misalignment. In panel d) there is only one yellow arrow on the top right corner because surface grid driven misalignment only occurs along a given layer. Note that: 1. in volumetric misalignment, only the misaligned ROI will include voxels from neighboring layers (panels b and c); 2. during volumetric misalignment, the training ROI can be shifted along large penetrating vessels that are orthogonal to the cortical surface (panel c); and 3. the distance between neighboring voxels within a layer is variable: it is equal to .8 mm (i.e. the length side of a voxel) if the voxels lie on the same plane, and 1.13 (i.e. the length of the diagonal of a voxel) if they do not. Within the surface grid driven regime therefore, a 1 voxel shift can lead to a misalignment that is greater than the nominal voxel resolution.

All misalignments were performed in Matlab. We imported the data in Matlab using the BVQX toolbox and used a number of in-house tools to misalign the test site relative to the training site.

#### 3.9.1 Volumetric misalignment

This approach consists of shifting the test ROI 0 to 5 voxels along the 3 axes (i.e. x, y and z) and 2 directions (i.e. positive and negative). Volumetric misalignment can be conceptualized by thinking of the training site as denoting the “origin” of a three-dimensional discrete Cartesian space, while the test site represents a point within said space whose coordinates differ from the origin along *one* of the 3 dimensions. Starting on the x axis, for example, we moved the test ROI 1 voxel in one direction (e.g. positive relative the origin). We then tested the accuracy of the classifiers, trained on the original ROI, with the misaligned site and stored that value. We then moved the test site again (by one additional voxel) along the same dimension and direction, and tested the accuracy of the classifiers’ models on this newly shifted ROI. We repeated this procedure for all other dimensions (i.e. y and z) and directions (i.e. positive and negative) to reach a total of 5 voxel shifts for all axes and directions. This approach led to a set of 6 (shifts i.e. 5 misaligned plus the original site) by 3 (dimensions) by 2 (directions) accuracy scores for each subject. We then computed the mean across dimensions and directions independently per voxel shift, leading to 6 accuracy scores (one per voxel shift). Importantly, misaligning the test site in volumetric space allows the inclusion of voxels belonging to neighboring layers in the shifted ROI.

#### 3.9.2 Surface grid driven misalignment

As the title suggests, we used the spatial coordinates of Laplace-based equipotential grids to spatially *guide* the misalignment of the test site in volume space. First, we identified the voxels belonging to a single cortical depth as indicated by the 3D coordinates of the Laplacian grids. We then removed up to 5 voxels per grid row at the medial most edge of the representation of the occluded quadrant. This procedure was implemented to allow shifting of the test site up to 5 voxels while still remaining within the boundaries of the representation of the occluded quadrant within each cortical depth, and ensuring that the number of voxels remained constant across misaligned ROIs. We trained the classifiers and tested their performance on the original ROI and on its shifted versions. As in the volume-based misalignment, the test ROI was shifted 1 to 5 voxels. Importantly, to ensure that the misaligned test ROI only included voxels within a given cortical depth, misalignment only occurred in one direction, specifically away from the medial portion of the occluded quadrant. This procedure ensured that displacement of the test ROI remained confined within the retinotopic representation of the occluded quadrant, to avoid potential confounds in our decoding results related to the inclusion of BOLD activity elicited by the stimulated portion of V1.

### 3.10 Univariate analyses

To test whether the 3 different images elicited different univariate BOLD amplitudes (defined as the mean activity across all voxels within a given cortical depth), we carried out the following statistical tests. We performed a 2 (signals) by 3 (images) by 6 (cortical depths) Linear Mixed Model (Matlab, The Mathworks Inc, 2014) with the mean BOLD response as a dependent variable. We combined the data from 4 runs and 4 participants (i.e. using 16 data points). Random variation across runs within each subject was accounted for by considering the subjects as random effects. This was implemented in Matlab using the following equation:

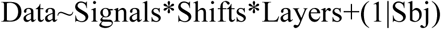

We estimated our fixed effects coefficients by means of maximum likelihood estimation. We computed hierarchical 95% bootstrap confidence intervals post hoc on significant main effects and interactions on the difference between the accuracies of the original ROI and its misaligned counterparts. We constructed a bootstrap distribution as follows: first, for a given subject, we computed the difference between the accuracies estimated on the unshifted ROI and those estimated for e.g. 1 voxel shift for all runs. For each bootstrap iteration, we then sampled with replacement these differences over the runs, computed the mean across the sampled runs and stored the value. We repeated this procedure for the remaining subjects, leading to 4 mean values (one per subject). We then sampled with replacement subjects and computed the mean, this time across subjects. We repeated this operation 2000 times. This procedure allowed us to construct a bootstrap distribution that is not limited by the factorial of N (number of subjects), as the same random sample of subjects would produce different bootstrap mean values due to the different runs sampled. We therefore computed the 95% confidence interval (Bonferroni corrected by adjusting the alpha threshold by the number of comparisons, in this case 5 – i.e. the number of voxels shifts) for these bootstrapped differences. Statistical significance was inferred when 95% bootstrap confidence interval did not overlap with 0. Only differences between the accuracies on the unshifted ROI and those on its displaced version were computed.

### 3.11 Univariate vs. Multivariate

To assess the relationship between univariate differences, univariate amplitudes and decoding accuracies across cortical depths, we performed a Spearman correlation analysis amongst these 5 measures (i.e. 3 decoding accuracies, univariate differences and univariate amplitudes) on the feed-forward signal for the original un-shifted ROI. To quantify the differences in activation across conditions, we computed univariate differences as follows: for each subject, run, image and trial, we computed the mean across the beta weights of each voxel (representing BOLD percent signal change, or PSC, amplitude). We then L2 normalized these values as follows: for each subject we put runs, images and trials in a vector and computed the L2 norm according to the following equation:

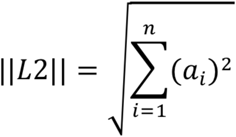

where n represents the length of vector *α* (i.e. 3 images x 4 runs x 6 trials). We divided the values of each subject by the L2 norm and computed the mean across trials. This procedure was implemented because we were interested in the differentials pattern across cortical depths, regardless of the raw differences in PSC BOLD amplitude across subjects. We then calculated the root square differences (RSD) of these normalized measures between each pair of images according to the following equation:

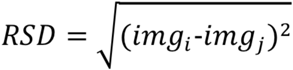

where img_i_ and img_j_ represent the L2 normalized activity elicited by 2 given images. We calculated the mean across the differences for all runs and image pairings. We choose RSD to quantify univariate differences in amplitude elicited by the 3 image stimuli because we wanted a measure that was insensitive to the sign of the difference, thus allowing averaging to quantify the mean difference across all image pairs.

We therefore computed Spearman correlation coefficients between: a) the 6 accuracy scores (i.e. one per cortical depth) for each of the tested classifiers and the 6 average univariate differences; b) the 6 accuracy scores for each of the tested classifiers and the 6 average univariate amplitudes; and c) the 6 average univariate amplitudes and the 6 average univariate differences.

Moreover, to assess the relationship between univariate amplitude and decoding accuracies and determine whether the former can explain the pattern of accuracies *across cortical depths and misalignment extents*, we carried out further correlational analyses. Independently, signal and layer, we calculated the mean across voxels, conditions, runs and trials for the original ROI and all its misaligned versions. We thus compared the resulting 6 (layers) by 6 (shifts) univariate amplitude matrices with the 6 by 6 matrices of the SVM accuracy scores, with the 6 by 6 matrices of the LDA accuracy scores, and with the 6 by 6 matrices of the NBC accuracy scores. To quantify the similarity between the decoding accuracies of all classifiers and univariate BOLD activation, we computed Spearman correlation coefficient between the accuracy scores and the univariate BOLD matrices.

### 3.12 Inferential statistic on decoding accuracies

For each of the 3 classifiers, to test the main effects and interactions between signals (feedforward and feedback), misalignment extents (1 to 6) and cortical depths (1 to 6), we performed a 2 (signals) by 6 (cortical depths) by 6 (voxel shifts) Linear Mixed Model (Matlab, The Mathworks Inc, 2014) with the accuracy scores as a dependent variable, as explained above. Moreover, we computed hierarchical 95% bootstrap confidence intervals post hoc on significant main effects and interactions as also described above.

## 4. Results

### 4.1 Univariate analysis

The 2 (signals) by 6 (cortical depths) by 3 (images) linear mixed model performed on BOLD amplitude averaged across all voxels within a given ROI cortical depth showed a significant main effect (p< .01) of signal (F(1,552)=14.554) driven by larger amplitudes elicited by feedforward stimulus conditions compared to feedback; and a significant (p< .05) interaction between signal and cortical depths (F(5,552)=2.342), driven by the fact that different cortical depths elicit significantly different amplitudes (increasing as we approach the pial surface) for the feedforward but not the feedback condition (see Figure 4A). Importantly, this analysis did not reveal significant differences between univariate BOLD elicited by the 3 different images. No additional significant main effects or interactions were observed.

**Figure 4.**
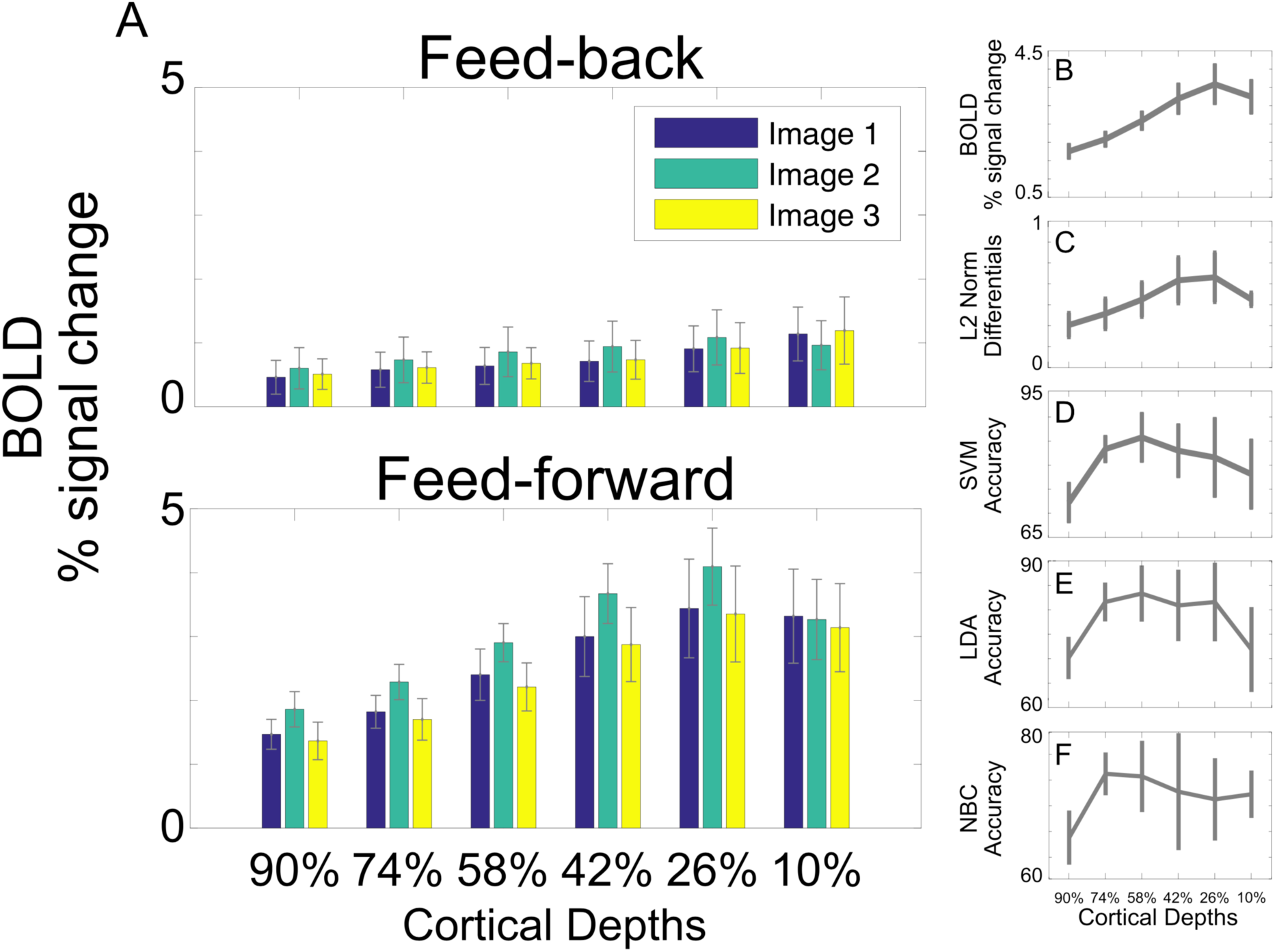
A) Univariate BOLD response for feed-back (top panel) and feed-forward (bottom panel) signals elicited by each image for all cortical depths. Errorbars represent the standard error across subjects. B) Mean univariate responses across images for the feed-forward signal. Errorbars represent the standard error across subjects. C) Mean univariate differences across images for the feed-forward signal. Errorbars represent the standard error across subjects. D) SVM decoding accuracy for the feed-forward signal. Errorbars represent the standard error across subjects. E) LDA decoding accuracy for the feed-forward signal. Errorbars represent the standard error across subjects. F) NBC decoding accuracy for the feed-forward signal. Errorbars represent the standard error across subjects.

We further report a strong positive Spearman correlation (rho(5) = .94; p=.0167) across cortical depths between feed-forward univariate averages (i.e. mean across images, trials and voxels), amplitudes and differences (figure 4B and 4C); no significant correlations (p>.05) were observed between feed-forward decoding accuracies (figure 4D, E and F) and either univariate amplitudes or average differences.

### 4.2 Decoding analysis

As previously reported (see Muckli et al., 2015), for the decoding analyses on the original un-shifted ROI, single-block classification was significant at each depth for each classifier and each subject (permutation tested at 5%; no corrections) during feedforward stimulation of V1. For the feedback condition (i.e. the occluded images), only the superficial outermost depth (10%) was significant in all four subjects and classifiers. The second-most outer depth (26%) was significant in three of four subjects for all classifiers, and the mid-depth (42%) was significant in two of four for SVM and LDA, and for 1 out of 4 for NBC. No subjects showed significant above chance decoding at the 58% mid-depth across all classifiers.

### 4.3 Misalignment on synthetic data

The results of the simulations are shown in figure 5.

**Figure 5.**
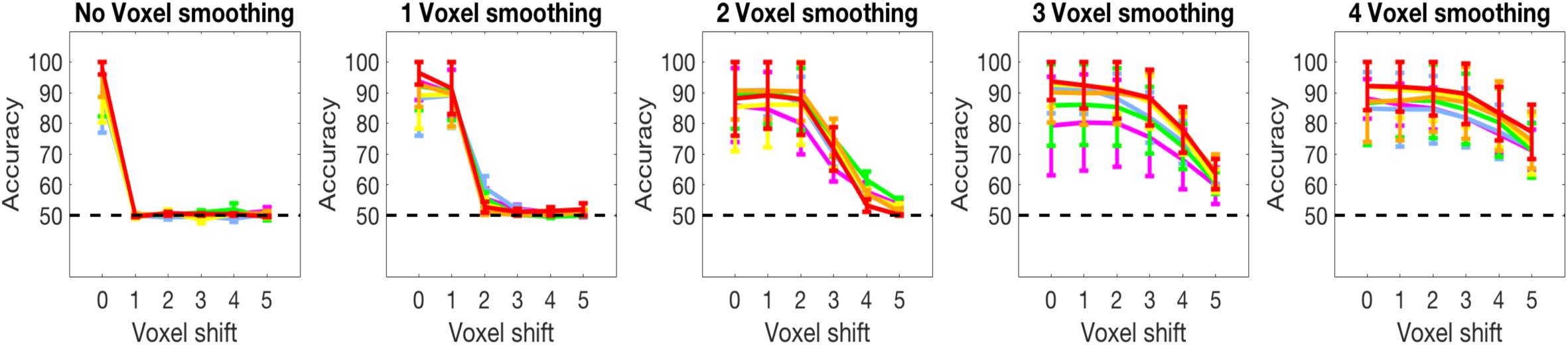
Impact of misalignment on multivoxel patterns with different spatial precision for data generated by injecting the synthetic multivoxel pattern to demeaned epi volume. Different colors portray different simulated cortical depths. Errorbars represent standard errors across simulated subjects.

We found that the resolution of the mutlivoxel pattern significantly modulates the impact of misalignment on SVM decoding accuracy: bonferroni corrected 95% bootstrap confidence intervals show that with no smoothing, a one voxel misalignment completely destroys the highly specific spatial structure of the patterns, rendering the model computed on the original ROI (i.e. training phase) redundant and causing decoding accuracy to immediately drop to baseline. Misaligning a mutlivoxel pattern smoothed with a 3D Gaussian kernel with a FWHM of one voxel required a 2 voxel shift to impair decoding accuracy; patterns smoothed with a FWHM of 2 voxels required a 3 voxel shift; FWHM of 3 voxels, a 4 voxel shift and a FWHM of 4 voxels required a 5 voxel misalignment to significantly impair decoding accuracy (figure 5).

### 4.4 Misalignment on real data

Artificial misalignment either led to a decrease or to no significant change in the accuracies of all classifiers for all layers, signals, and misalignment scenarios (see sections below).

While we carried out a fully parameterized linear mixed model, we were specifically interested in the 3-way interaction between cortical depths, signals, and voxel shifts. In the following sections, F values for all significant main effects and interactions captured by the model are reported; however, a more in-depth discussion of these numbers is outside the scope of this work. We focus on the interactions amongst cortical depths, signal and voxel shifts, and the related post-hoc bootstrap tests.

#### 4.4.1 Volumetric misalignment

The 2 (signals) by 6 (cortical depths) by 6 (voxel shifts) linear mixed model carried out on SVM accuracy showed significant main effects (p< .01) of signals (F(1,1080)=289.04) and depths (F(5,1080)=24.806) as well as significant (p< .01) interactions between signals and cortical depths (F(5,1080)=12.995), signals and voxel shifts (F(5,1080)=37.749), cortical depths and voxel shifts (F(25,1080)=3.2061) and signals, cortical depths and shifts (F(25,1080)=1.7484, p< .05).

The same analysis carried out on LDA accuracies, showed significant main effects (p< .01) of signals (F(1,1080)=241.92) and depths (F(5,1080)=24.69) as well as significant (p< .01) interactions between signals and cortical depths (F(5,1080)=16.067), signals and voxel shifts (F(5,1080)=30.983), cortical depths and voxel shifts (F(25,1080)=3.252) and signals, cortical depths and shifts (F(25,1080)=2.1793, p< .01).

The linear mixed model for NBC also showed significant main effects (p< .01) of signal (F(1,1080)=175.92) and depths (F(5,1080)=15.715) as well as significant (p< .01) interactions between signal and cortical depths (F(5,1080)=8.756), signal and voxel shifts (F(5,1080)=19.865), and signals, cortical depths and voxel shifts (F(25,1080)=1.8607).

For the feed-forward signal, post-hoc 95% bootstrap confidence interval (btCI) revealed that a one voxel shift produces a significant decrease in decoding accuracies of all classifiers for all cortical depths (Figure 7). Conversely, for the feedback signal, 95% btCI showed that for the outermost cortical depth (i.e. 10%) a 2 voxel shift is required before observing a significant decrease in decoding accuracies for all classifiers; for LDA, a 2 voxel shift also led to a significant drop in decoding accuracy for the second and third outermost depths (i.e. 26% and 42%) (Figure 7).

Spearman coefficients were computed to quantify the similarity between decoding accuracies and univariate BOLD activation (Figure 6), by correlating decoding accuracies for all classifiers and univariate BOLD responses at all misalignment extents independently per cortical depth and signal. We observed no significant correlations (q>.05 FDR corrected).

**Figure 6.**
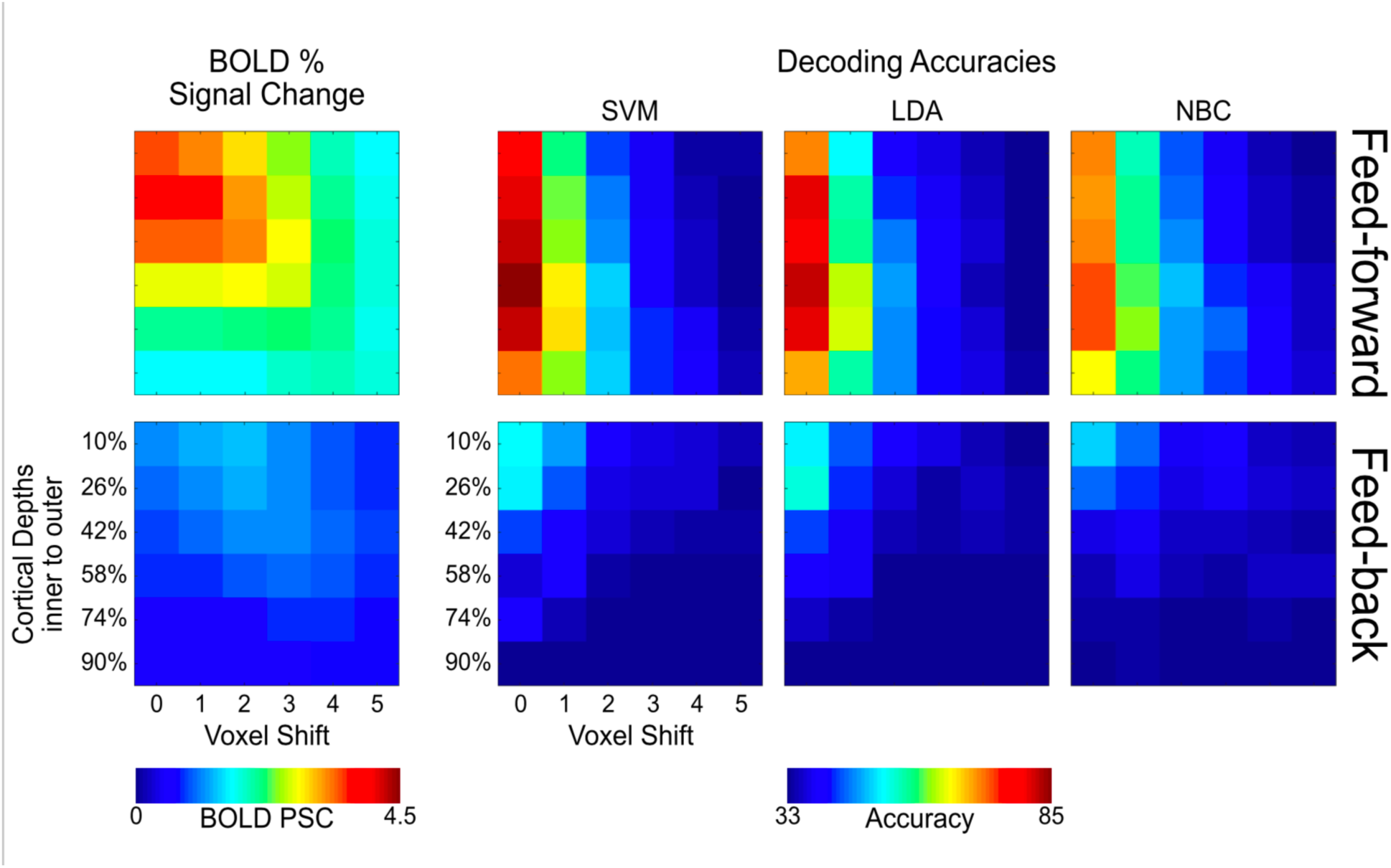
Volumetric univariate BOLD amplitude vs. volumetric SVM, LDA and NBC accuracies across misalignment extents (x-axis) and cortical depths (y-axis) for the feed-forward (top panels) and feed-back (bottom panels) signals.

**Figure 7.**
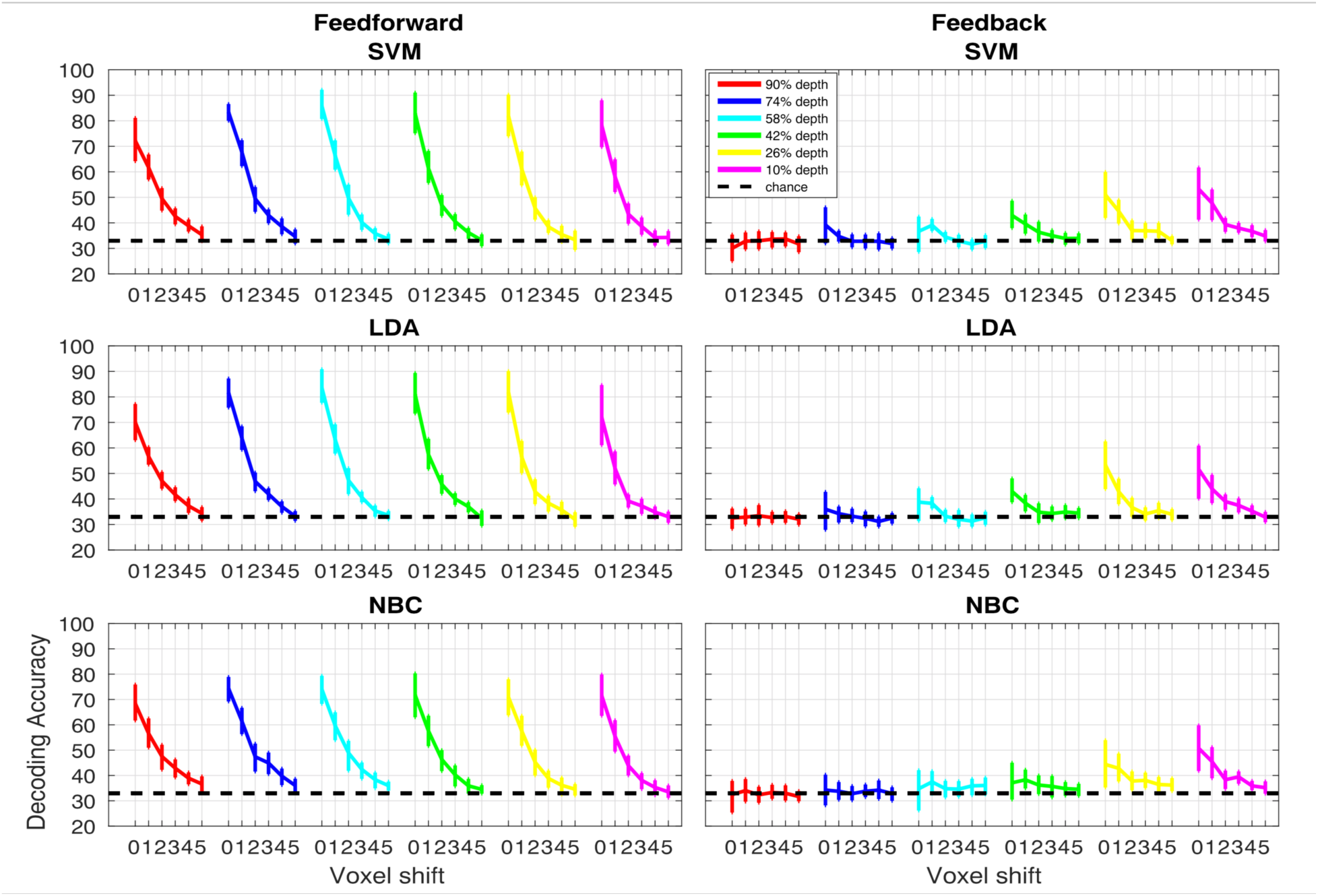
Volumetric misalignment. Decoding accuracy as a function of voxel misalignment for the feed-forward (left panels) and feed-back (right panels) signals for SVM, LDA and NBC. Different colors represent different cortical depths as indicated in the figure legend. Error bars show the 95% bootstrap confidence interval.

In table 1, we report the total number of voxels independently per subject and cortical depth, after performing the contrast t_target_ > t_surround_ ∩ t_target_ > 0 ∩ t_full-stimulus_ > t_occluded-stimulus_.

**Table 1.**
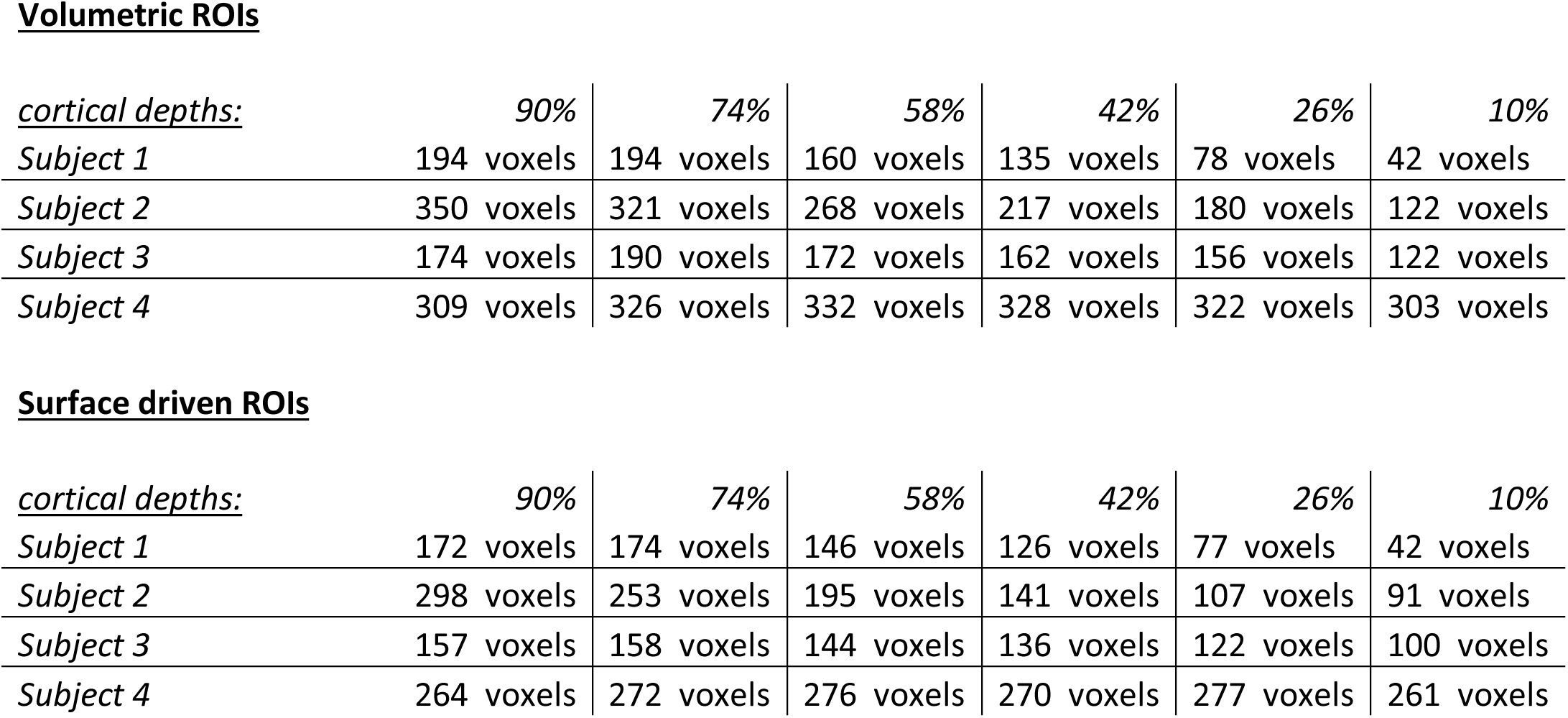
Table 1 reports the number of voxels included in each cortical depth for the ROIs used for the volumetric and surface driven misalignment.

**Volumetric ROIs**
| *cortical depths:* | 90% | 74% | 58% | 42% | 26% | 10% |
| --- | --- | --- | --- | --- | --- | --- |
| *Subject 1* | 194 voxels | 194 voxels | 160 voxels | 135 voxels | 78 voxels | 42 voxels |
| *Subject 2* | 350 voxels | 321 voxels | 268 voxels | 217 voxels | 180 voxels | 122 voxels |
| *Subject 3* | 174 voxels | 190 voxels | 172 voxels | 162 voxels | 156 voxels | 122 voxels |
| *Subject 4* | 309 voxels | 326 voxels | 332 voxels | 328 voxels | 322 voxels | 303 voxels |

| *cortical depths:* | 90% | 74% | 58% | 42% | 26% | 10% |
| --- | --- | --- | --- | --- | --- | --- |
| *Subject 1* | 172 voxels | 174 voxels | 146 voxels | 126 voxels | 77 voxels | 42 voxels |
| *Subject 2* | 298 voxels | 253 voxels | 195 voxels | 141 voxels | 107 voxels | 91 voxels |
| *Subject 3* | 157 voxels | 158 voxels | 144 voxels | 136 voxels | 122 voxels | 100 voxels |
| *Subject 4* | 264 voxels | 272 voxels | 276 voxels | 270 voxels | 277 voxels | 261 voxels |

#### 4.4.2 Surface grid driven misalignment

The 2 (signals) by 6 (cortical depths) by 6 (voxel shifts) linear mixed model carried out on SVM accuracy showed significant main effects (p< .01) of signal (F(1,1080)=131.1) and depths (F(5,1080)=10.291), as well as significant (p< .01) interactions between signal and cortical depths (F(5,1080)=7.635), signal and voxel shifts (F(5,1080)=18.302), cortical depth and voxel shift (F(25,1080)=1.989), and signal, cortical depths and shift (F(25,1080)=1.886, p< .01).

The same analysis carried out on LDA accuracies, showed significant main effects (p< .01) of signal (F(1,1080)=98.074) and depths (F(5,1080)=8.606) as well as significant (p< .01) interactions between signal and cortical depths (F(5,1080)=9.585), signal and voxel shifts (F(5,1080)=12.651), cortical depth and voxel shift (F(25,1080)=2.539) and signal, cortical depths and shift (F(25,1080)=2.513, p< .01).

The linear mixed model for NBC also showed significant main effects (p< .01) of signal (F(1,1080)=112.17) and depths (F(5,1080)=6.854) as well as significant (p< .01) interactions between signal and cortical depths (F(5,1080)=6.665), signal and voxel shifts (F(5,1080)=14.653) and signal, cortical depths and shift (F(25,1080)=1.663, p< .05).

For the feed-forward signal, post-hoc 95% bootstrap confidence interval (btCI) revealed that one voxel shift produced a significant decrease in decoding accuracies of all classifiers for all cortical depths (Figure 9). For the feedback signal, 95% btCI showed that for the 2 outermost cortical depths only (i.e. 10% and 26%), a 1 voxel shift produced a significant decrease in SVM accuracies (Figure 9); for LDA accuracy, a 1 voxel shift led to a significant decrease for the 2^nd^ outermost depth only (i.e. 26%), while for the outermost depth, a 2 voxel shift was necessary to significantly impair decoding accuracy; for NBC accuracy, a 2 voxel shift led to a significant decrease for the outermost depth only (i.e. 10%).

We further carried out Spearman correlations to quantify the similarity between the MVPA decoding accuracy and univariate BOLD activation (Figure 8). Spearman coefficients, computed independently per cortical depth and signal by correlating decoding accuracies and univariate BOLD response at all misalignment extents, revealed no significant correlations (q>.05 FDR corrected).

**Figure 8.**
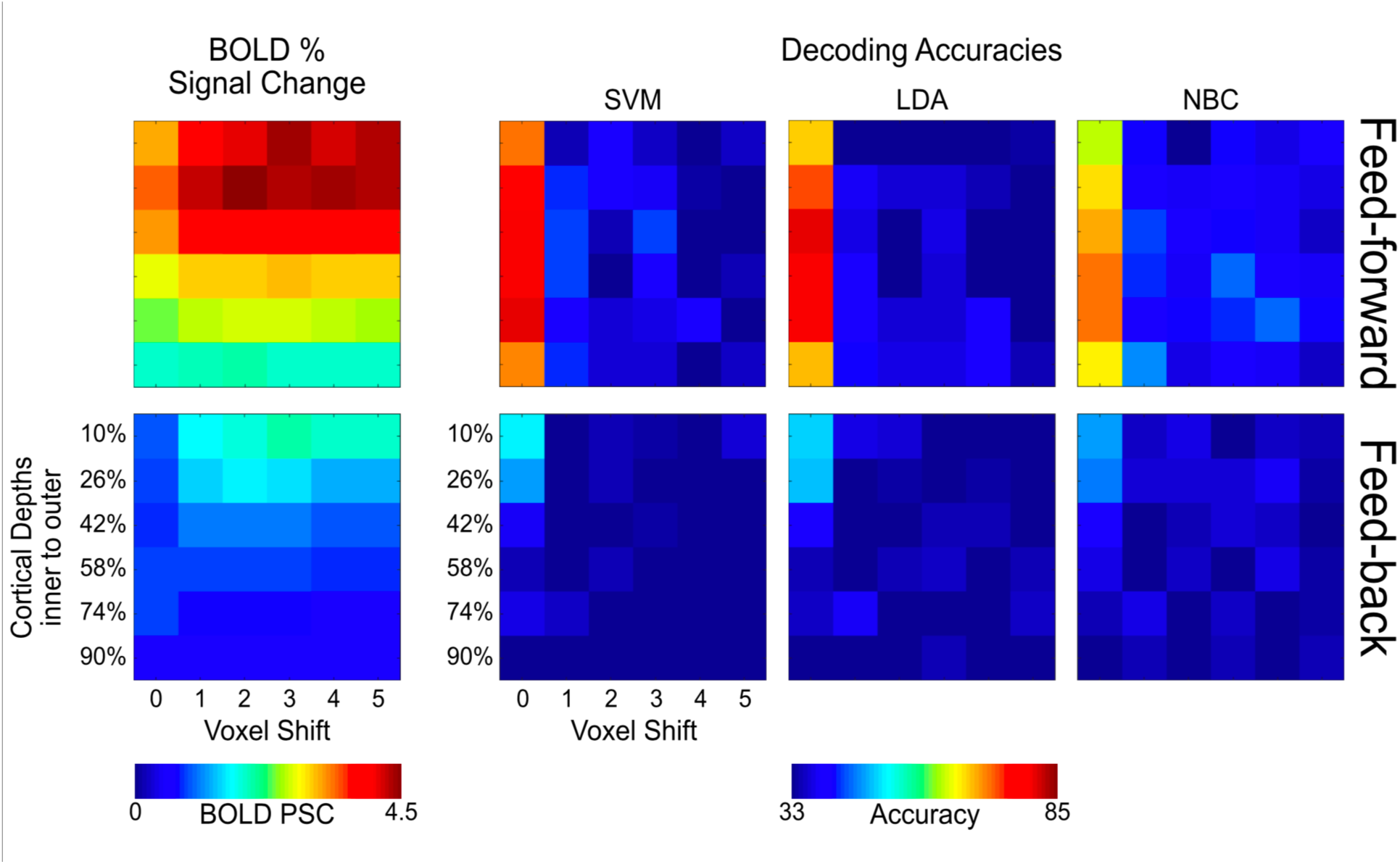
Iso-surface grid driven univariate BOLD amplitude vs. volumetric SVM, LDA and NBC accuracies across misalignment extents (x-axis) and cortical depths (y-axis) for the feed-forward (top panels) and feed-back (bottom panels) signals.

The total number of voxels after performing the contrast t_target_ > t_surround_ ∩ t_target_ > 0 ∩ t_full-stimulus_ > t_occluded-stimulus_ independently per subject and cortical depth are reported in table 1.

## 5. Discussion

The ability to exploit the sub-millimeter resolution achievable with UF fMRI is critical for advancing cortical depth dependent functional investigations in humans. This is particularly true for the widely used GE BOLD contrast, which has high signal-to-noise ratio, but limited spatial acuity. We measured whether MVPA is capable of relying on stimulus specific fine scale responses.

With previously collected data from Muckli et al. (2015), we parcellated the cortical sheet into 6 equally spaced depths, ranging from 10% to 90% distance from the pial surface. We analyzed feed-back and feed-forward signals triggered by images of natural scenes in V1 (Muckli et al., 2015) independently per cortical depth.

To assess whether MVPA relies on fine scale stimulus specific responses we systematically misaligned voxels between the training and test ROI. We trained decoding algorithms (linear Support Vector Machine (SVM), Linear Discriminant Analysis (LDA) and Naïve Bayes Classifier (NBC)) on a given cortical depth and tested their performances on a ROI that was misaligned anywhere from 0 to 5 voxels relative to the training site. This approach allowed us to assess whether information decoded with MVPA is at least as precise as the nominal resolution of single voxels. We hypothesized that a negligible decrease in decoding accuracy following the spatial offset of the test ROI relative to the training ROI would suggest that the exact correspondence of spatial structures is not necessary to achieve the highest decoding accuracy, indicating that the multivoxel pattern is blurred and/or the information decoded exists at a scale coarser than the tested offset. Conversely, a significant decrease in decoding accuracy following a 1 voxel misalignment would indicate that the exact correspondence of spatial structures is necessary to achieve the highest decoding accuracy, suggesting that the scale of responses exploited by multivoxel decoders and associated with the tested stimuli is at least as precise as the nominal resolution of single voxels (here 0.8 mm isotropic). The results of our simulation on synthetic data support our hypothesis, suggesting that the spatial scale of the multivoxel pattern of BOLD activation modulates the drop in decoding accuracy and, therefore, that our method does measure stimulus specific scale of BOLD responses.

When we applied the misalignment approach to real data we found that as little as a one voxel misalignment led to a significant decrease in decoding accuracy across all cortical depths. We argue that multivoxel activity patterns carry a substantial amount of spatially precise information at the nominal resolution of single voxels. Our result suggests that multivoxel decoding can enhance the relevance of voxels that are not corrupted by reduced specificity, thereby increasing the sensitivity of the multivoxel approach to fine-grained spatial responses.

### 5.1 Distinguishing different sources of spatial resolution

We first briefly distinguish the related, yet different sources of spatial resolution of interest. We identify two sources of spatial resolution: 1) those related to acquisition, including the nominal resolution of BOLD images and the functional resolution of single voxels (for example, as measured by PSF); and 2) those related to post-processing or analytical operations, such as the resolution of the multivoxel pattern exploited by MVPA.

Our focus is to measure the resolution exploited by multivoxel decoders, and to understand whether MVPA profits from the nominal resolution afforded by UF fMRI (here .8 mm isotropic). The question was partly motivated by the observation that point spread measurements of GE BOLD recordings suggest that the point spread function of GE BOLD responses is above the millimeter range (e.g. Shmuel et al., 2007; see “Relation to BOLD PSF” below). These reports not only challenge the feasibility of imaging human cortical layers and columns, but also question the usefulness of recording BOLD images with nominal sub-millimeter resolution. But is the resolution of the multivoxel pattern of BOLD activity exploited by multivariate decoding also impacted by the limited precision of BOLD measurements? Here we show that this is not the case, and argue that multivariate decoders exploit fine-grained information contained in multivoxel patterns of activity. Our finding that a 1 voxel shift leads to a significant impairment in decoding accuracy indicates that: 1) the resolution of the multivoxel pattern of BOLD responses exploited by MVPA is finer than that of single functional voxels as measured by PSF; 2) while BOLD PSF results render the investigation of the mesoscale organization of the human cortex in the sub-millimeter range challenging at best, alternative analytical strategies, such as MVPA or differential maps (e.g. Yacoub et al., 2007), seem to permit this submillimeter scale; 3) unlike univariate BOLD amplitude, which is limited in spatial acuity, using MVPA we can fully exploit the nominal resolution of functional voxels (here .8 mm isotropic); and 4) decoding of complex image stimuli, containing a mixture of low and high spatial frequencies, relies both on fine and coarse patterns of multivoxel activity. These arguments are now discussed in more detail.

### 5.2 Relation to BOLD PSF

As suggested by studies measuring the GE BOLD point spread function (PSF) of single voxels at 7 T (e.g. Shmuel et al., 2007; Chaimow et al., 2018; Parkes et al., 2005), the spatial specificity of individual functional voxels extends beyond the millimeter range, despite their nominal resolution. The GE BOLD PSF for a single condition in humans has been reported to have an upper limit of 2 mm when avoiding large vessels (Shmuel et al., 2007; but see Chaimow et al., 2018). If we take this measurement at face value, shifting the multivoxel pattern of activity by 0.8 mm (i.e. one voxel) should have a negligible impact on decoding accuracy. This is because, despite the 0.8 mm nominal resolution, neighboring voxels should yield highly correlated signals, effectively decreasing the functional resolution of the voxel population. Our results demonstrate, however, that a one-voxel misalignment significantly impacts decoding accuracy, indicating that MVPA decoding operates at a resolution that is at least as precise as the nominal voxel size (i.e. .8 mm isotropic), thus fully exploiting the increase in spatial resolution affordable with GE sequences at 7 T, and further motivates the need for sub-millimeter GE recording. These results are somewhat in line with those reported by Chaimow et al. (2018), who estimate a PSF of 1.02 mm for GE BOLD. This however is not to say that the BOLD information content is at the same spatial precision, as a number of factors, for example vascular heterogeneity, may be contributing to the pattern of results observed here.

### 5.3 Pulse sequence limitations in high-resolution fMRI

It is well known that spatial specificity of the BOLD response is strongly modulated by large draining vessels (Bandettini et al., 1994; Boxerman et al., 1995; Constable et al., 1993; Duong et al., 2002; Duong et al., 2003; Duyn et al., 1994; Frahm et al., 1994; Kim et al., 1994; Lai et al., 1993; Lee et al., 1995; Menon et al., 1995; Menon et al., 1993; Segerbath et al., 1994; Shmuel et al; 2007; Song et al., 1996; Ugurbil et al., 1999; Uludag et al., 2009; Yacoub et al., 2007; Yacoub et al., 2005; Yacoub et al., 2003; Yacoub et al., 2001). This modulation is, to an extent, dependent on the type of contrast used. Spin echo (SE) based acquisitions are less susceptible to large veins and thus less affected by venous artifacts compared to gradient echo (GE) sequences (Yacoub et al., 2007; Yacoub et al., 2005; Yacoub et al., 2003) and therefore seemingly represent the ideal choice to maximize BOLD spatial specificity. However, SE based acquisitions are limited in terms of coverage, SNR, and CNR. In this respect, GE acquisitions represent an appealing choice for high resolution fMRI at high fields. However, the high sensitivity of GE BOLD data are related to the impact of large draining veins (Duyn et al., 1994; Frahm et al., 1994; Kim et al., 1994; Lai et al., 1993; Lee et al., 1995; Menon et al., 1993; Segebarth et al., 1994; Shmuel et al; 2007; Song et al., 1996; Ugurbil et al., 1999; Uludag et al., 2009; Yacoub et al., 2007; Yacoub et al., 2005; Yacoub et al., 2003; Yacoub et al., 2001). Venous BOLD signal, demonstrated in both humans (Krings et al., 1999; Lee et al., 1995) and animals (Keilholz et al., 2006; Silva et al., 2007), produces a larger response compared to that of tissue (Gati et al., 1997; Yacoub et al., 2001). Moreover, the vascular architecture of the human cortex, characterized by a higher concentration of large draining veins in proximity of the pial surface, represents an additional challenge in imaging layers using GE sequences. The typical GE laminar BOLD response is characterized by a ramping-like profile with larger amplitude, SNR, and CNR in outer layers (e.g. Goense and Logothetis, 2006; Goense et al., 2007; Ress et al. 2007; Polimeni et al., 2010; Koopmans et al., 2010; Koopmans et al., 2011). Such a bias makes investigating depth dependent functional responses challenging, especially when only evaluating mean univariate amplitudes. At the multivoxel level, however, we found that decoding accuracy in human primary visual cortex does not co-vary with univariate BOLD amplitude and amplitude differences. We observed that decoding accuracy, at least for the feed-forward signal, peaks over the mid layers, while the univariate BOLD response and its differences peak in the outer depths. To further evaluate this uncoupling, we directly compared the relationship between univariate amplitude and MVPA decoding accuracy by correlating these 2 metrics for all cortical depths and misalignment extents (Figures 6 and 8). We observed no significant correlation across layers and signals. Differential mapping can help to avoid the effects of larger vessels and overcome the lack of high spatial specificity typically observed in GE data (e.g. Cheng et al., 2001; Dechent and Frahm, 2000; Goodyear and Menon, 2001; Menon et al., 1997; Yacoub et al., 2007). We therefore wanted to assess whether the impact of large draining vessels on mean *univariate* differences is comparable to that observed on mean univariate BOLD amplitudes. Large vessels, prominently distributed on the pial surface, are known to increase BOLD response, giving rise to the widely observed ramping pattern of BOLD amplitudes across cortical depths (i.e. larger at the outer compared to the inner depths). As indicated by a strong positive correlation between univaritate differences and average amplitudes, we report a comparable, albeit shallower, profile of the average univaritate differences magnitudes and BOLD amplitudes across cortical depths, both peaking in the outer depths. Conversely, no correlation was observed between mean univariate amplitudes or their differences and MVPA decoding accuracy, with the latter peaking over the mid cortical depths. Moreover, as mentioned, large vessels compromise the spatial specificity of GE BOLD. Yet, for the feedforward signal, a 1 voxel misalignment on decoding accuracy always leads to a significant decrease in decoding accuracy across all depths, in spite of the diverse distribution of vein across the cortical ribbon. These results indicate a comparable spatial precision of multivoxel patterns of activity across depths.

Taken alone, the layer profile observed for decoding accuracy (peaking in the mid, rather than outer depths) may reflect hypersensitivity of MVPA to physiological vascular noise, prominent in outer depths due to the high concentration of large vessels. Such noise may limit the performance of MVPA in outer depths. While this still represents a plausible scenario, the decoding layer profile, together with the comparable impact of misalignment across depths, suggest that, unlike univariate analyses, MVPA is relatively less contaminated by the effects of large vessels. It is worth noting that univariate differences as computed here, although comparable, are not the same as differential mapping. The latter does not imply spatial averaging, preserving the spatial structure of the BOLD response. Differential mapping, like MVPA, thus exploits the pattern of activation, as opposed to its average, and could therefore be less susceptible to the effect of large vessels (Yacoub et al., 2007). While the debate regarding the optimal sequence choice to study layers and columns is not yet settled, we argue that clever post-processing and analytical tools represent a viable path to overcome sequence-based resolution and specificity issues.

### 5.4 Univariate amplitude vs. multivoxel decoding accuracy

The feed-forward pattern of decoding accuracy observed here, peaking over the mid cortical depths, together with the observation that a 1 voxel shift equally impacts MVPA decoding accuracy across depths, demonstrate that univariate BOLD amplitude does not (in this case) modulate mulitvoxel decoding accuracy. This result is in line with a previous report showing that, while macroscopic vessels can carry neuronal-specific information, their contribution to mulitvoxel decoding accuracy may be redundant (Shmuel et al., 2010; Yao et al., 2017).

Moreover, we report that standard univariate contrast, defined as the average BOLD amplitude across all voxels within a given cortical depth, does not show significant differences between the amplitudes elicited by different images for either the feed-forward or the feed-back conditions (Figure 4). We argue that this result stems from the fact that different images elicit differently retinotopically distributed maps of activation, and that these differences are obscured following voxels averaging (for a review of the differences between multivariate and univariate analysis see Davis et al., 2014).

### 5.5 Finer vs. Coarser scale information in MVPA

Importantly, while misaligning the test ROI by 1 voxel negatively impacts decoding accuracy, as demonstrated by the volumetric approach, significant above chance decoding can still be achieved with as many as three voxel shifts. This result indicates that the multivoxel pattern of activity carries both coarser- and finer-grained information.

Several approaches have estimated whether MVPA relies on stimulus specific fine scale responses, ranging from smoothing (Op de Beeck, 2010) or spatially filtering (Swisher et al., 2010) the activation maps in order to degrade the resolution of the information, to shifting the slice positioning by 1 mm (i.e. half a voxel) during the acquisition (Freeman et al. 2013). While spatial filtering is a generally useful strategy, we argue that, within the context of this work, it does not represent a straightforward advantage over misalignment for a number of reasons. Low pass filtering, for example, increases SNR and decreases run to run variation (e.g. Alakorkko et al., 2017), potentially boosting cross validated decoding accuracy (see Figure S3). Additionally, smoothing introduces artificial, spurious correlations across voxels (e.g. Korhonen et al., 2017) and this effect and its impact on cross-validated decoding accuracy is difficult to quantify. Moreover, in light of the observed reliance of decoding on both finer and coarser spatial information, together with the afore mentioned low-pass filtering induced increase in SNR, down-sampling the input pattern will not necessarily lead to a drop in accuracy, even if multivoxel decoding does rely on spatially precise patterns of activation. The complex and poorly understood interplay of these forces would therefore render down-sampling related modulations on cross-validated multivoxel decoding accuracy difficult to interpret.

The approach implemented by Freeman’s et al. (2013), although different as the misalignment occurred in the acquisition phase, is conceptually comparable to the approach adopted here. Freeman et al. (2013), however, did not find the 1 mm misalignment to significantly decrease decoding accuracy. A number of differences between Freeman’s study and the current one may explain the apparent discrepancy. First, Freeman et al. used 3 T fMRI with a nominal voxel resolution of 2 mm isotropic. Not only are the magnitude of the detected signal changes and the spatial scale different compared to this 7 T study, but the vascular contributions are as well (Ugurbil, 2016). Further, the data analyzed here come from a block design experiment, while Freeman et al. used a fast temporal encoding paradigm. Fast temporal-encoding paradigms have been shown to artificially enhance the impact of coarse global maps on MVPA (Pratte et al., 2016). Moreover, Freeman et al. used highly controlled low-level visual stimuli (i.e. spiral gratings), while here we used images of visual scenes, rich in both low and high spatial frequencies. As such, the findings reported here provide intuitions into the sensitivity of multivoxel decoding to high and low spatial frequency information when the input stimuli contain both low and high spatial frequencies, which is not guaranteed to be the same as when low or high spatial frequency stimuli are presented in isolation. A recent study by Alink et al. (2017) implemented an analysis strategy comparable to the one adopted here to assess the spatial scale of information exploited by MVPA during orientation decoding in V1. They spatially offset the testing relative to the training set by 1, 2, 4, and 6 mm and measured the impact on decoding accuracy. Alink et al. found that a 1 mm shift significantly impaired orientation decoding accuracy. These results are somewhat comparable to the ones reported here. It is worth pointing out though that a number of important differences exist between Alink’s and our work. Their data were recorded at 3 T with 2 mm isotropic voxels and their functional images were interpolated offline to 1 mm isotropic before the misalignment was performed. Moreover, Alink et al., like Freeman, used a tailored, low-level stimulus set to directly test orientation decoding in V1. All these studies, regardless of the results, debate whether MVPA can retrieve finer-grained information than what is observable via conventional univariate analyses. Here, instead, we use our approach to test whether the finer spatial scale of information observed with a multivoxel pattern of activity persists when the functional voxels are sampled with sub-millimeter isotropic voxels and where the BOLD sensitivity and PSF are expected to be the limiting factors in the ability to observe such information. In light of the limited spatial acuity of GE functional voxels, the usefulness of the super-high resolution achievable with GE sequences can be questionable. One may ask if there is an advantage in acquiring sub-millimeter images, if voxels’ amplitudes are highly correlated, effectively decreasing the functional resolution of the data. We claim that by using MVPA, we can fully exploit sub-millimeter resolution of GE BOLD fMRI at UF and that MVPA therefore represent a viable strategy to study the mesoscale organization of the human cortex.

It is worth making one more consideration regarding the performance of the different classifiers implemented here. All classifiers led to very similar results, with only minor differences. The most obvious difference is that NBC led to overall lower decoding accuracy compared to SVM and LDA. This is probably due to the fact that NBC assumes orthogonality amongst the variables within a class (i.e. zero off-diagonal covariance) and it is thus less flexible than SVM and LDA. However, the impact of misalignment on decoding accuracies led to consistent results for all 3 classifiers, demonstrating that the results are likely to be a property of the resolution of the multivoxel pattern exploited by MVPA rather than specific to a classification algorithm.

### 5.6 Differences between volume and surface based misalignments

We implemented artificial misalignment using 2 approaches: 1) volume-based misalignment, in which the test ROI was misaligned along all dimensions and directions, thus allowing trespassing into neighboring cortical depths; and 2) surface grid-driven misalignment, in which misalignment occurred only within a given cortical depth (Figure 4). While both approaches show that a single voxel misalignment leads to a significant decrease in decoding accuracy, we observe differences between the 2 techniques. Volumetric misalignment yields a smoother function of accuracy across space or voxel shifts (Figures 7 and 9), taking at least 3 shifts before decoding accuracy drops to chance level. Conversely, grid-driven misalignment yields a sharper function, where, across layers and signals, one voxel shift leads to a drastic decrease in decoding accuracy. We hypothesize that the difference between the approaches is related to: 1) a sparser voxel sampling for the grid compared to the volumetric-based misalignment; and 2) by constraining the spatial offset of the training ROI within a given depth we effectively avoid large penetrating draining vessels perpendicular to the cortical surface. The sparser sampling is a direct outcome of constraining the misalignment within a layer. As depicted in figure 4, for the volumetric misalignment the distance between neighboring voxels is invariant and equates to the width of a voxel (in this case 0.8 mm), whereas for the surface grid driven approach, the distance between 2 voxels can be greater in at least 2 scenarios: 1) when 2 voxels that are adjacent to one another in volume space belong to different depths, therefore introducing a one voxel “gap” between 2 neighboring voxels within a cortical depth; and 2) when 2 adjacent voxels belonging to the same depth do not lie on the same plane, rendering the distance between them to be equal to the square root of the sum of their squared edge length (i.e. the voxel’s diagonal). Sparser sampling leads to a greater impact of one voxel shift on decoding accuracies because when only considering activity within a given layer, voxels are spatially farther apart and thus less correlated. Spatially offsetting the test relative to the training ROI would therefore lead to greater misalignment for the grid-driven compared to the volumetric approach. Moreover, given that during surface grid-driven misalignment the spatial offset of the training relative to the test ROI only occurs *within* a depth (see figure 4) and therefore tangentially to the cortical surface, this approach is less affected by the effect of large penetrating vessels that contribute to blurring single voxels’ functional specificity. We further observe that volumetric misalignment has a lesser impact on decoding the feedback compared to the feedforward signal, requiring a two-voxel shift before significantly decreasing feedback accuracy. As previously argued (Muckli et al., 2015), this observation could suggest that the feed-back signals from higher-level regions with larger receptive fields carry information that is more abstract and therefore spatially coarser than its feed-forward counterpart. While this claim is supported by a recent study directly investigating the contribution of spatial frequencies to feedback signal (Revina et al., 2018), as indicated by the size of the error bars, this finding could simply be related to noisier signal rather than an apparent coarser resolution. Moreover, as previously mentioned, we observed that decoding accuracy is differentially modulated by misalignment for feedback and feedforward signals. Unlike feedforward, for the feedback signal decoding accuracy shows no significant decrease following a one voxel shift. The differential modulatory impact of misalignment on feedback and feedforward signals represents a further indication that our results are not a mere property of decoding analyses (or else we would expect a comparable pattern of accuracy decrease across signals, see also figure S1 in the supplementary section).

**Figure 9.**
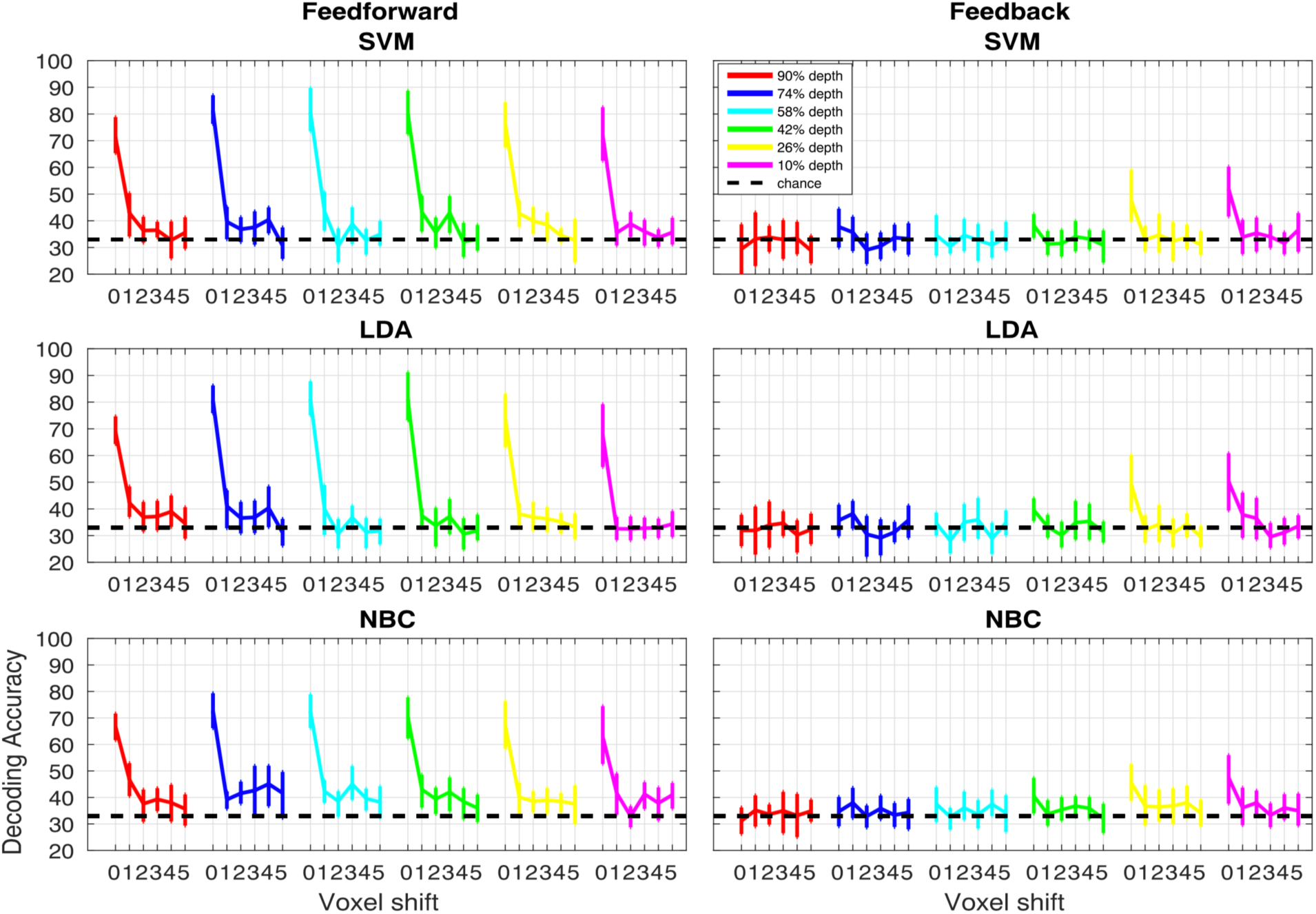
Iso-surface grid driven misalignment. Decoding accuracy as a function of voxel misalignment for the feed-forward (left panels) and feed-back (right panels) signals for SVM, LDA and NBC. Different colors represent different cortical depths, as indicated in the figure legend. Error bars show the 95% bootstrap confidence interval.

## 6. Conclusion

We showed that the multivoxel pattern of activity exploited by MVPA decoding carries information about the visual experimental condition on a finer scale, one that here is not visible with a mean univariate analysis. This finding is promising for future fMRI studies of cortical layers and columns, as it indicates that, when the spatial precision of mean univariate amplitude is corrupted by macroscopic biases such as, for example, large draining vessels or blurring in the phase encoding direction, MVPA can potentially circumvent sensitivity and specificity limits of the GE BOLD signal. While there are several pulse sequence variants that could reduce the large vessel biases present in high field GE BOLD data, such as SE or VASO, they are costly in terms of efficiency and sensitivity. As an alternative, intelligent analysis strategies provide benefits in enhancing the spatial precision of the information in fMRI signals.

## Acknowledgements

We thank Dr. Junpeng Lao for the useful discussions on analytical procedures, and Yulia Revina for help with reference data.

## Funding

This project has received funding from the European Union’s Horizon 2020 Framework Program for Research and Innovation under the Specific Grant Agreement No. 720270 and 785907 (Human Brain Project SGA1 and SGA2) and European Research Council (ERC StG 2012_311751-‘‘Brain reading of contextual feedback and predictions’’ to LM).

