## Supplemental Material for "Multivoxel Pattern of Blood Oxygen Level Dependent Activity can be sensitive to stimulus specific fine scale responses"

### Supplementary section.

**Large vessels, SVM weights, voxel selection and misaligned decoding accuracy.**

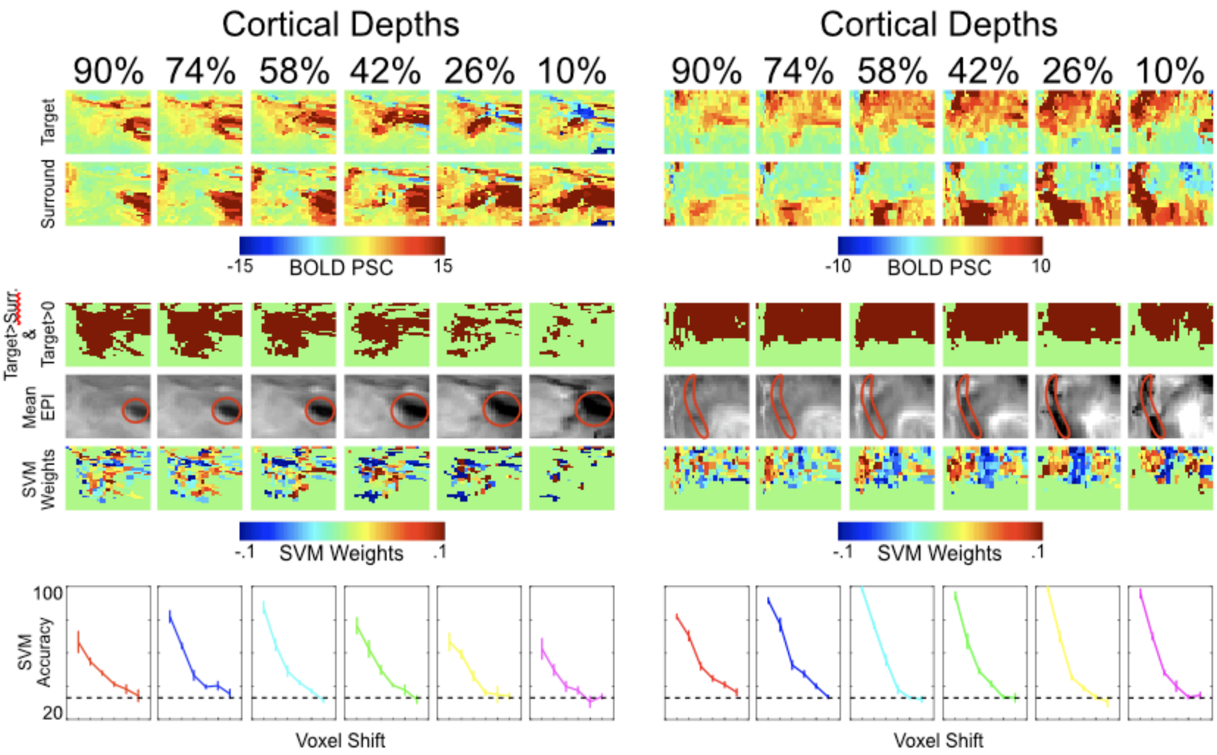


**Figure S1.** Example of 2 subjects. The top row shows the beta weights elicited by the target condition. The second row shows the beta weights elicited by the surround condition. The third row depicts the patch of voxels included in the analysis as determined by the contrast: target > surround ∩ target > 0. The fourth row shows the mean epi. Within the red contour we can see a large draining vessel (in dark), prominent in proximity of the pial surface. The fifth row shows the average SVM weights. The sixth row portrays the SVM accuracy across misaligment extent (voxel shift). Error bars represent standard error across cross-validated folds.

While large draining vessels are known to compromise BOLD signals, especially for GE data, our data show that cortical vasculature alone cannot explain the misalignment results. In fact, as argued in the main body of this work, we believe that veins have a minimal (or at least a lesser) impact on MVPA. Cortical vascular architecture has a defined structure, with a higher concentration of large draining vessels in the outer pial surface. This organization leads to the known ramping BOLD amplitude pattern with GE sequences (observed here as well). If the drop in accuracy following misalignment was a mere vascular effect, we should observe differential modulations of misalignment across cortical depth. This is not the case as, regardless of the cortical depth, one voxel shift leads to a significant decrease of SVM accuracy.

As evident from figure S1:

1. SVM weights do not look like a venous map. This observation, together with the differential impact of misalignment on feedforward and feedback signals, and the comparable impact of feedforward misalignment across cortical depths, indicates that venous cortical architecture alone cannot explain the results presented here.

2. Voxel selection (i.e. target > surround ∩ target > 0) can result in a more or less contiguous patch, depending, amongst other things, on the presence of large draining vessels. For example, for the subject portrayed on the left hand side (figure S1), the presence of a large vein (highlighted by the red circle) within

the retinotpic representation of the bottom right hand quadrant of the visual field (i.e. the “target” region), compromises the contiguity of the selected voxels’ patch (especially when approaching the pial surface). On the other hand, however, the subject shown on the right hand side has a large vein (red contour) outside the retinotopic representation of the target area. Consequently, the resulting selected voxels’ patch is in fact continuous. Crucially, the impact of misalignment on decoding accuracy remains comparable (i.e. one-voxel shift leads to a significant impairment of SVM accuracy) in spite of the difference in the sparseness of the voxel selection, both across subjects and and across cortical depths within subjects (see for example the subject portrayed on the left). In sum, the sparseness of the activated voxels differs substantially not only between the subject portrayed on the left that that portrayed on the right, but also clearly differs across cortical depths for the subject on the left. Yet, in spite of these differences, misaligning the test ROI by one voxels always leads to a significant decrease in decoding accuracy.

3. The voxel selection criterion implemented here (i.e. target > surround ∩ target > 0) tends to render less likely the inclusion of such voxels whose signal is compromised by the vicinity of large vessels. Target versus surround voxel selection will result in selecting only voxels that are entirely in the target area and that are precisely tuned (i.e. consistently showing a greater response to the target relative to the surround condition). When we are close to vessels the tuning should be less accurate and target and surround signals should become more mixed and hence excluded from analysis.

#### Equidistant vs. Equivolume.

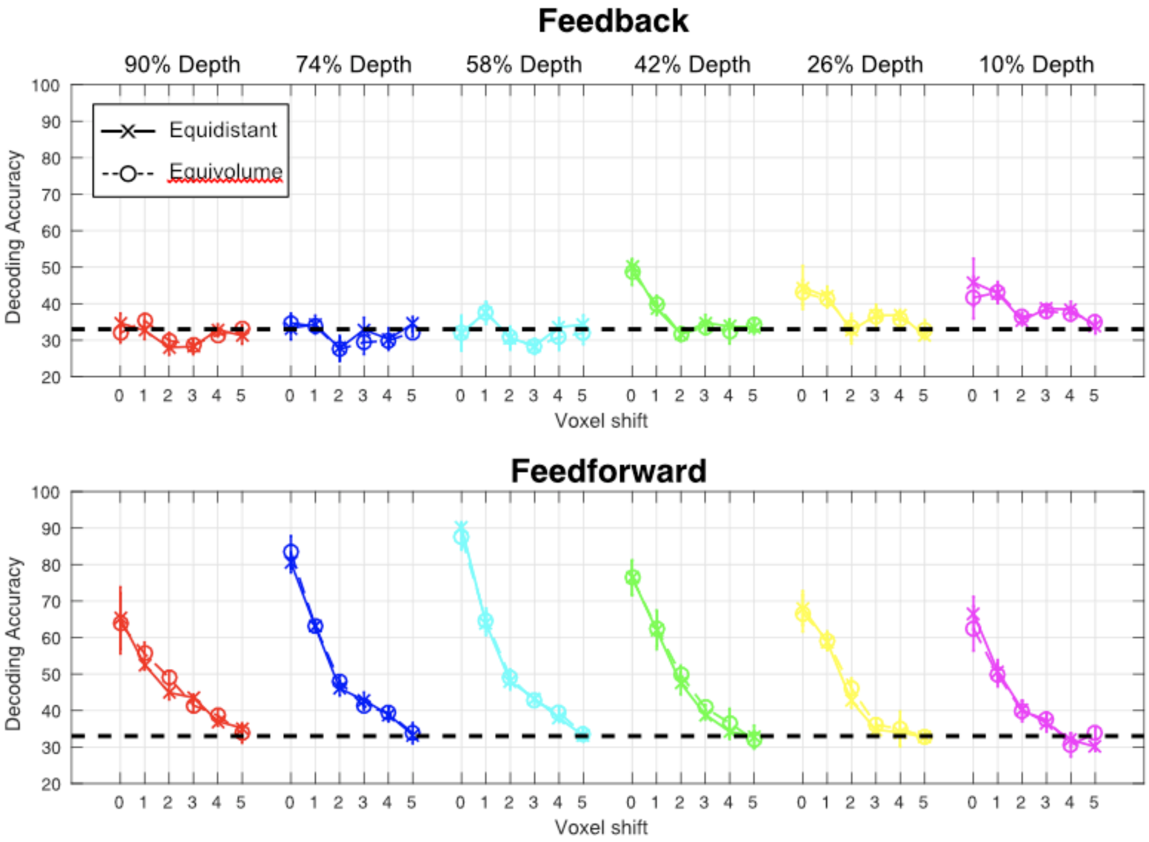


**Figure S2.** Equidistant vs equivolume cortical depth sampling. Example of direct comparison for 1 representative subject of the impact of misalignment on equidistant and equivolume cortical depth sampling. Error bars represent standard error across cross-validated folds.

Equivolume segmentations tends to be regarded as being more accurate (at least anatomically). However, the choice of equidistant cortical depth stems from 3 main arguments:

*1.* Recent evidence shows that, at the resolution of .8 mm isotropic, difference between equidistant and equivolume cortical depths sampling is minimal (e.g. see Kemper et al., 2018)
*2.* We wanted to ensure consistency with Muckli et al. (2015)
*3.* In our specific case, we observe no differences between the 2 cortical depth sampling approaches (see figure S2 for an example)

In light of these consideration we opted for the equidistant cortical depth sampling, which has the additional advantage of being is more straightforward and making fewer assumptions than its equivolume counterpart.

#### Misalignment vs Smoothing

While some of the arguments listed below have already been mentioned in the main manuscript, we re-state them here for ease of read/

A number of previous studies have implemented a down-sampling approach to estimate the spatial scale of information in fMRI (Gardumi et al. 2016, Alink et al. 2017, Mandelkow et al. 2017). We do not believe, at least within the specific context of this work, that spatially filtering the data would represent a straightforward advantage over the approach implemented here for a number of reasons:

1. Low pass filtering increases SNR (e.g. Alakorkko et al., 2017).
   - Low pass filtering will increase SNR and decrease run to run variation, therefore potentially increasing cross validated decoding accuracy.

2. Low pass filtering introduces artificial, spurious correlations across voxels (e.g. Korhonen et al., 2017).
   - This effect and its impact on decoding accuracy is difficult to quantify.

3. We use complex stimuli portraying natural scenes.
   - The stimuli used here contain both high and low spatial frequency. Accordingly, our results indicate that decoding accuracy is underpinned by both high and low spatial frequency patterns. Filtering out high spatial frequency from the input pattern will therefore not necessarily lead to a drop in decoding accuracy in light of the reliance of SVM decoding on lower spatial frequency patterns and the increase in SNR related to the filtering operation (see point 1).

4. Low pass filtering is known to shift the peak of activation and the activation pattern itself (e.g. Ridgway et al., 2012).
   - smoothing can turn negative voxels into positive ones, rendering the decoder's weights associated with those voxels potentially inaccurate and thus leading to a decrease in decoding accuracy, regardless of the spatial resolution of the multivoxel pattern.
   - Moreover, potential shifts in peak activation are especially problematic for the feedback conditions, as it may lead to including activity coming from the stimulated portion of V1 in the retinotopic representation of the occluded quadrant.

In light of these observations, we believe that the effects of low pass filtering on SVM accuracy will be difficult to interpret. While a comparison with down-sampled data would be interesting, given the complexities of this comparison, it would be difficult to fully address within the confinements of this work. Nonetheless, for completeness, over the next pages we show the impact of spatial smoothing on SVM accuracy.
Based on the premise that variation in BOLD activation across runs and trials to the same image can be regarded as noise-related, we try to quantify this SNR by Pearson correlating the multivoxel pattern of BOLD response elicited by the training data with that elicited by the testing data, across trials and runs. Unsurprisingly, the average correlation coefficient across voxels, computed amongst all runs and trials, is lowest for the innermost depth (see figure S5). Importantly, smoothing significantly increased the mean correlational values for all depths, but most prominently for the innermost depth (see figure S5).
This increase in SNR, loosely defined here as consistency across trials and runs, will lead to an increase in cross-validated decoding accuracy. Concomitantly, if it is true that decoding accuracy also relies on fine-grained spatial information, it would be fair to assume that smoothing should reduce SVM classification performance. However, the increase in SNR after smoothing may obscure any “information loss”, resulting in no changes or even in increase in decoding accuracy, ultimately making it difficult to interpret the cross-validated SVM performance. This would seem to be what is happening in our specific case, where the observed decrease in decoding accuracy following 1 voxel

volumetric misalignment is in the order of approximately 10-20% (see figure 5); while, Pearson coefficients increase by approximately 200-300% following a 1 voxel smoothing (see figure S5). More specifically, while decoding accuracy seems to always increase when downsampling (i.e. smoothing) to a simulated voxel size of 1.6 isotropic voxels, such an increase is only significant for the 2 innermost depths, where we also observe the largest increase in trials consistency across runs (as quantified by Pearson correlation).

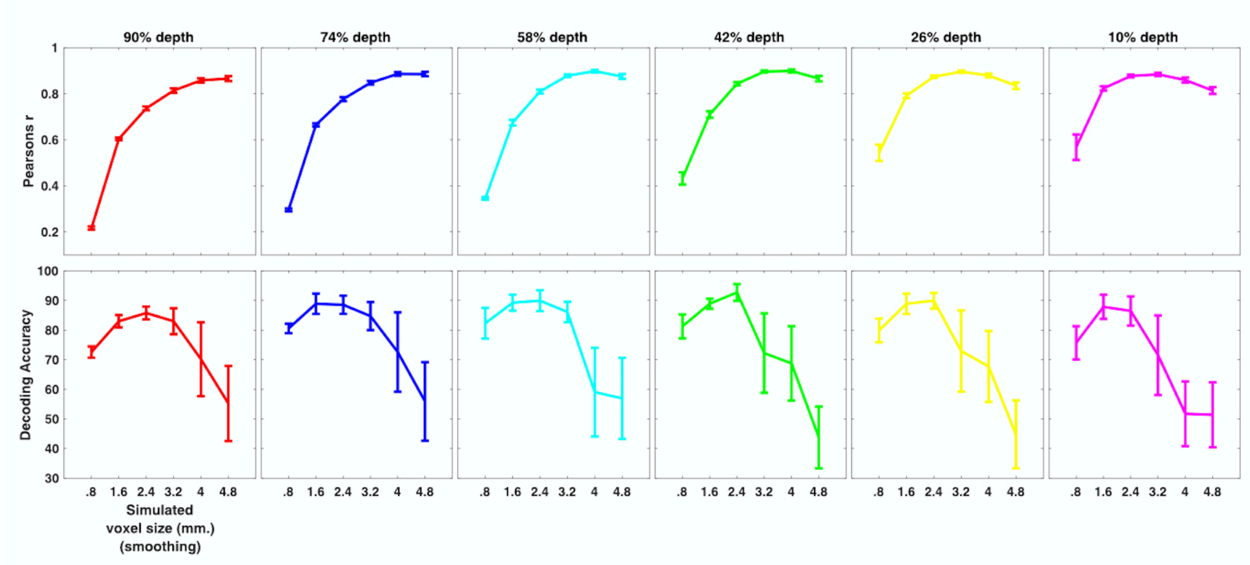


**Figure S3.** Impact of smoothing on SNR (top row) and decoding accuracy (bottom row). Errorbars represent the standard errors across subjects.
